# Attachment/detachment of cortical myosin regulates cell junction exchange during cell rearrangement

**DOI:** 10.1101/2022.08.12.503738

**Authors:** Keisuke Ikawa, Shuji Ishihara, Yoichiro Tamori, Kaoru Sugimura

## Abstract

Epithelial cells remodel cell adhesion and change their neighbors to shape a tissue. This cell rearrangement proceeds in three steps: the shrinkage of a junction, exchange of junctions, and elongation of the newly generated junction. Herein, by combining live imaging and physical modeling, we showed that the formation of myosin-II (myo-II) cables around the cell vertices underlies the exchange of junctions. The local and transient detachment of myo-II from the cell cortex is coupled with the junction shrinkage and elongation via an interplay between the LIM domain-containing protein Jub and the tricellular junction protein M6. Furthermore, we developed a mechanical model based on the wetting theory and clarified the way by which the physical properties of myo-II cables are integrated with the junction geometry to induce the transition between the attached and detached states and support the unidirectionality of cell rearrangement. Collectively, the present study elucidates the orchestration of geometry, mechanics and signaling for exchanging junctions.

## Introduction

Cell rearrangement plays a fundamental role in shaping a tissue and developing multicellular patterns^1–5^. Cell rearrangement proceeds in three steps: the shrinkage of a junction, exchange of junctions around the cell vertex (four-way or higher-folded), and elongation of the newly generated junction. The molecular mechanisms underlying junction shrinkage and elongation have been well characterized. At the apical level, medial actomyosin flow and junctional recruitment of myosin-II (myo-II) lead to the generation of forces that shrink and elongate a junction^2–6^. At the basolateral level, actin-rich protrusions deform the cell membrane^7^. In sharp contrast, little is known regarding how epithelial cells exchange junctions during cell rearrangement.

Molecules comprising the junctional structure play essential roles in the development of epithelial tissue, where the bicellular junction (bCJ) including the adherence junction (AJ) adheres cells, and the tricellular junction (TCJ) seals the tissue at the cell vertex^2,8,9^. During cell rearrangement, as the bCJ dynamically changes its length, the position of the TCJ changes. The junction deformation and vertex displacement are determined by the balance between the constriction force generated by myo-II and the adhesive force generated by cell adhesion molecules such as E-cadherin^2–6^. The mechanical force balance is tuned by actin-AJ linkers, which are responsible for the linkage between actomyosin and E-cadherin, and TCJ proteins, which are a class of proteins comprising the cell vertex structures, to maintain cell adhesion during the junction shrinkage and control the speed of junction elongation^10–15^. The localization and activity of these molecules need to be coordinated such that cell adhesion is weakened specifically at short junctions around the cell vertex and such that *de novo* formation of cell adhesion is initiated between the correct pair of cells. The mechanism underlying the coordination of cell adhesion and the regulatory molecules during the junction exchange is unclear.

Recent studies from others and us identified a cytoskeleton structure that could be critical in the junction exchange^10,12,16^. We showed that myo-II transiently forms rectangle-shaped cables around the cell vertices at the AJ plane in *Drosophila* wing epithelium (Fig. 1a, b; hereafter called rsMC (<u>r</u>ectangle-<u>s</u>haped <u>m</u>yo-II <u>c</u>ables))^16^. Such local detachment of myo-II from short junctions has been reported in other epithelial tissues^10,12^. The rsMC may represent a temporally loosened junctional structure, which facilitates the reconnection of junctions. However, the molecular and physical basis of rsMC dynamics and function have not been studied extensively.

**Fig. 1.**
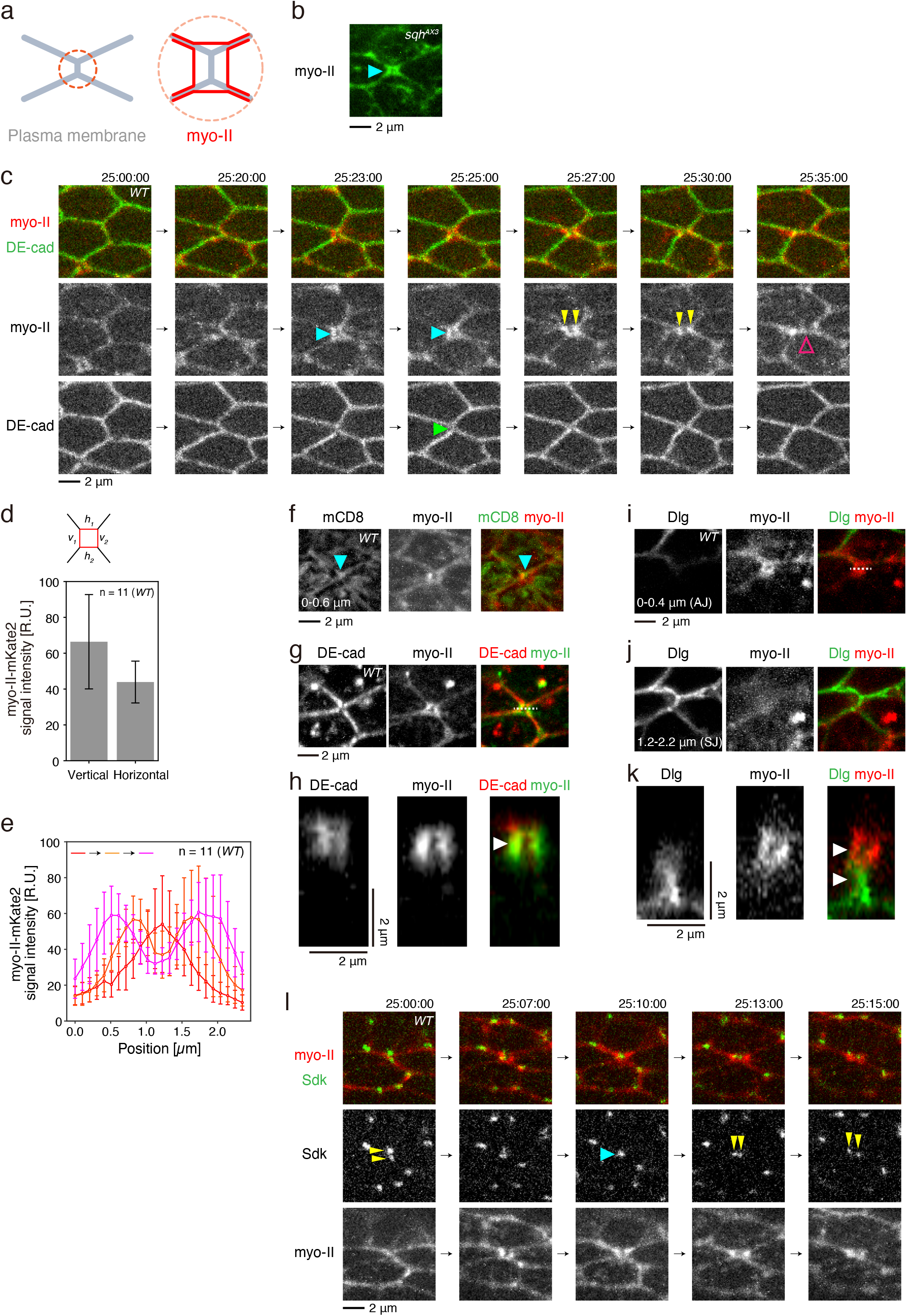
myo-II is transiently detached from the AJ during junction exchange. (a) Schematics of the cellular structure analyzed in this study. An orange circle in the left panel indicates the region shown in the right panel. Gray and red lines indicate the plasma membrane and cortical myo-II. (b) Image of myo-II-GFP at 24 h APF in a null mutant background of *spaghetti-squash* (*sqh*), which encodes a *Drosophila* gene of myosin regulatory light chain. Blue arrowhead points to the rsMC. (c) Time-lapse images of myo-II-mKate2 (red in upper panels, gray in middle panels) and DE-cad-GFP (green in upper panels, gray in bottom panels) during cell rearrangement in the *Drosophila* pupal wing. Blue, yellow, and magenta arrowheads indicate the detachment of myo-II from the AJ, the separation of the myo-II signals, and the formation of the PD junction, respectively. A green arrowhead points to the DE-cad signal inside the rsMC. (d, e) Quantification of the myo-II-mKate2 signal intensity based on time-lapse data captured at 25–26.5 h APF at 25 °C. (d) The myo-II-mKate2 signal intensity along the vertical (AP) or horizontal (PD) edges (*v_1_*, *v_2_*, *h_1_*, and *h_2_* in a schematic shown in the top) of rsMC at the time point when the rsMC was formed. (e) The myo-II-mKate2 signal intensity around the rsMC is measured along the line ROI, which aligns along the horizontal (PD) axis. Line colors represent the timing of measurement as shown in the upper left corner, where red, orange, and magenta lines correspond to 3 min before the rsMC formation, the time of rsMC formation, and 1 min before the rsMC fission, respectively. (f) Images of mCD8-GFP (gray in left panels, green in right panels) and myo-II-mKate2 (gray in middle panels, red in right panels) at 24 h APF. Arrowheads point to the mCD8-GFP signal inside the rsMC. (g, h) Images of an AJ marker, DE-cad-mTagRFP (gray in left panels, red in right panels) and myo-II-GFP (gray in middle panels, green in right panels) at 24 h APF. The vertical section along the dashed line in (g) is shown in (h). (i–k) Images of an SJ marker, Dlg-GFP (gray in left panels, green in right panels) and myo-II-mKate2 (gray in middle panels, red in right panels) at 24 h APF. (i, j) Top-view along the different z-planes indicated at the bottom. (k) Side-view along the dashed line in (i). (l) Time-lapse images of Sdk-YFP (green in upper panels, gray in middle panels) and myo-II-mKate2 (red in upper panels, gray in bottom panels) during cell rearrangement. Arrowheads point to the Sdk-YFP signal at reconnecting TCJs. Mann-Whitney *U* test: Vertical vs. Horizontal, P < 0.001 (d). The number of rsMCs (d) and the number of ROIs (e) examined are indicated. Data are presented as the mean ± s.d. (d, e). Scale bars: 2 μm (b, c, f–i, k, l).

Here, we sought to identify the mechanism by which apical junctions are exchanged during cell rearrangement in the *Drosophila* wing epithelium. The findings demonstrate that the precise control of rsMC formation supports the exchange of junctions and the stabilization of newly generated junctions. The rsMC formation is coupled with the junction shrinkage and elongation via an interplay between the LIM domain-containing protein Jub and the tricellular junction protein M6. A mechanical model based on the wetting theory can explain the junction-length-dependent transition between the detached and attached states of cortical myo-II and the unidirectionality of cell rearrangement. The coupling between junction geometry, mechanics, and signaling identified in the present study may function in other aspects of morphogenesis, as all morphogenetic events are associated with spatio-temporal changes in these three aspects.

## Results

### Local and transient formation of rectangle-shaped myo-II cables during junction exchange

To address the mechanism by which epithelial cells exchange junctions, we analyzed the distribution of molecules during cell rearrangement in the *Drosophila* wing epithelium (see Supplementary Fig. 1a, b for schematics of the wing). Following the shrinkage of anterior-posterior (AP) junction, myo-II detached from the junction (Fig. 1b, blue arrowheads in Fig. 1c, Supplementary Video 1). We designated the subcellular structure as the rsMC (<u>r</u>ectangle-<u>s</u>haped <u>m</u>yo-II <u>c</u>ables). DE-cadherin (DE-cad) became more diffusively distributed around the junctions (green arrowhead in Fig. 1c), and the plasma membrane marker mCD8-GFP was localized inside the rsMC (blue arrowhead in Fig. 1f). These data indicate that a junction structure was present inside the rsMC. As shown in Fig. 1g–k, a rsMC was formed at the AJ labelled by DE-cad, apical to the septate junction (SJ) labeled by Discs large (Dlg).

rsMC was a transient structure with an average duration time of 4.3 ± 4.1 min (n = 69). The myo-II signal intensity was, on average, higher along the vertical (AP) edges of the rsMC (Fig. 1d; we excluded medial pools of actomyosin from our analysis because they are not densely accumulated in the pupal wing). The separation of two bright spots of myo-II (yellow arrowheads in Fig. 1c) proceeded concomitantly with the decrease in the myo-II signal intensity on the horizontal edges of rsMC (Fig. 1e). Eventually, myo-II became more uniformly distributed along the newly formed proximal-distal (PD) junction, indicating the reattachment of myo-II cables to the junction (magenta arrowheads in Fig. 1c).

As cortical myosin cables dynamically change their shape during junction exchange, reorganization of other cellular structures occurs. We found that bright dots of Sidekick (Sdk)^10–12^, a tricellular AJ (tAJ) protein, apposed inside the rsMC, were then separated along the horizontal direction as a new junction was formed (Fig. 1l). This observation suggests that the disassembly and reassembly of TCJs occur at the AJ level during junction exchange. To probe actin cytoskeleton, we clonally expressed a F-actin marker, Life-Act-GFP. Prior to the separation of myo-II cables, actin protrusions invaded the rsMC at the AJ, whereas no significant actin-rich protrusions were detected at the basolateral side of the cells (Supplementary Fig. 1k, l, Supplementary Video 2). Collectively, these data showed that in the wing, the junction exchange is accompanied by the formation of the orderly structure comprising myo-II, the actin cytoskeleton and junction components at the AJ.

### rsMC formation is regulated by actin-AJ linkers including Jub

Next, we searched for molecules underlying rsMC dynamics. We identified actin-AJ linkers that strengthen the linkage between actomyosin and AJ, including the LIM domain-containing protein Jub/Ajuba and PDZ protein Canoe (Cno)/Afadin^14,15,17–21^, as important regulators of the process. Our time-lapse analysis detected pleiotropic phenotypes in *jub* RNAi cells: (i) the precocious formation of rsMCs from longer junctions, which resulted in the enlargement of the rsMC, (ii) prolonged duration of the rsMC, and (iii) failure in the reattachment of myo-II to the newly formed PD junction, which resulted in the destabilization of new junction and subsequent reverse reconnection of junctions in ~30% of events (Fig. 2a, c–e, Supplementary Fig. 2a, b, Supplementary Fig. 3c, Supplementary Video 3). *jub* RNAi cells retained a plasma membrane marker mCD8-GFP in the enlarged rsMC (Fig. 2f), suggesting that *jub* RNAi phenotypes were not due to the defect of the plasma membrane. *cno* RNAi and *Da-catenin (Dα-cat*) RNAi, but not *vinculin* (*vinc*) RNAi, altered the rsMC dynamics and resulted in the failure in junction exchange, similar to *jub* RNAi (Fig. 2b–e, Supplementary Fig. 2c–e, Supplementary Fig. 3c, Supplementary Video 3). The signal intensity of myo-II and DE-cad hardly changed on *vinc* RNAi, whereas *jub* RNAi decreased the DE-cad level and *cno* and *Da-cat* RNAi increased the myo-II level (Supplementary Fig. 3a, b). The absence of rsMC malformation in *vinc* RNAi cells was consistent with the hierarchy in co-dependence of actin-AJ linkers in their subcellular distribution (Supplementary Fig. 4 and its legends). The correlation coefficient between the rsMC area and average area of cells surrounding the rsMC was 0.04 (Fig. 2g and Supplementary Fig. 1i), which strongly suggests that the enlargement of rsMC was not the secondary consequence of the change in cell size.

**Fig. 2.**
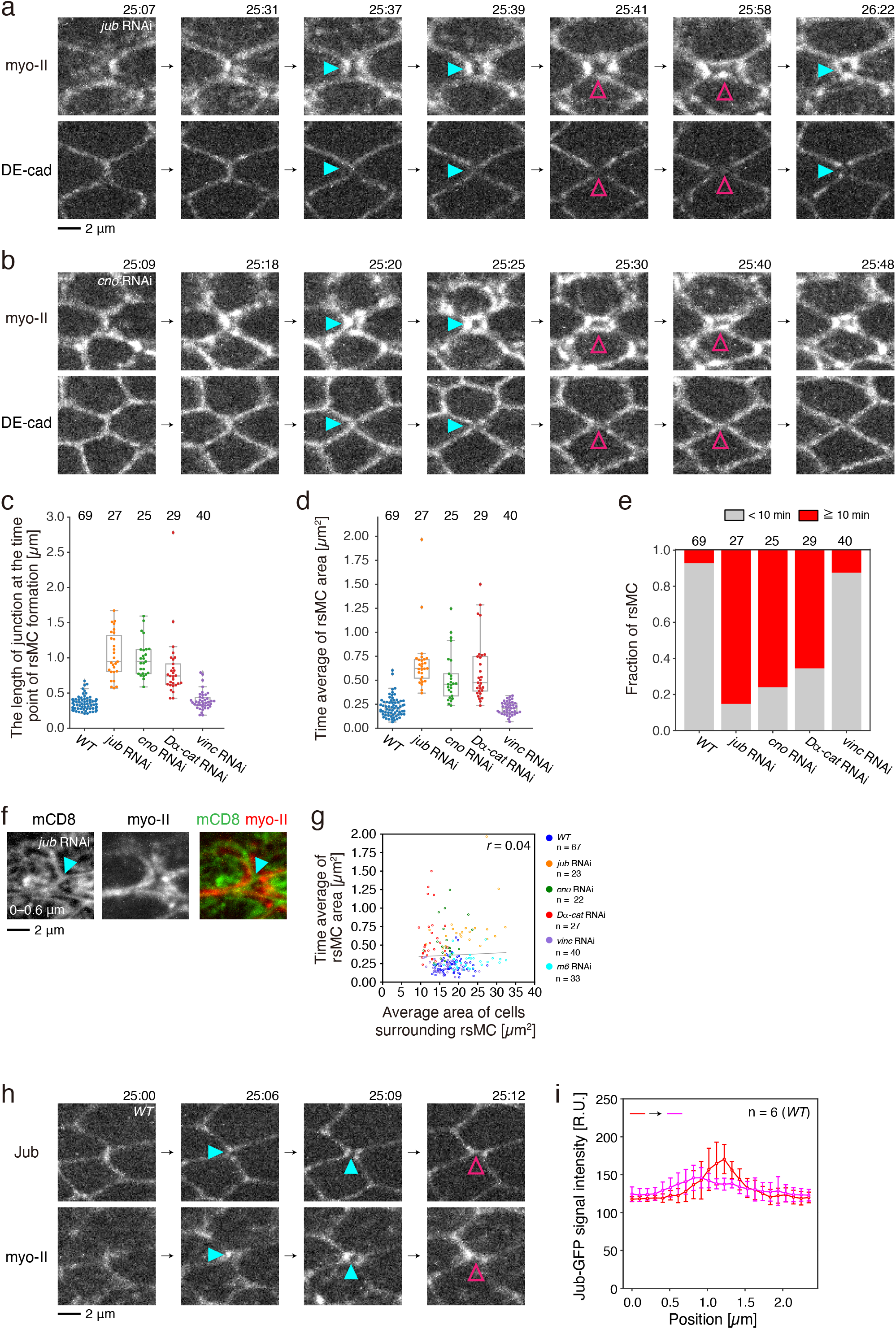
The Jub-GFP signal is transiently weakened during junction exchange. (a, b) Time-lapse images of myo-II-mKate2 (upper) and DE-cad-GFP (lower) with the indicated genotypes (a, *jub* RNAi and b, *cno* RNAi). Blue and magenta arrowheads point to the rsMC. (c–e) Quantification of RNAi phenotypes. Junction length at the time point of rsMC formation (c), the temporal average of rsMC area (d), and the fraction of rsMC time duration (e) for each genotype based on time-lapse data captured at 25–26.5 h APF. In (e), rsMC time duration is defined as the length of time between the formation and fission of the rsMC. (f) Images of mCD8-GFP (gray in left panel, green in right panel) and myo-II-mKate2 (gray in middle panel, red in right panel) at 24 h APF in *jub* RNAi cells. Arrowheads point to the mCD8-GFP signal inside the rsMC. (g) The correlation between the time average of rsMC area and the average area of four cells surrounding the rsMC with the indicated genotypes (blue: *WT*, orange: *jub* RNAi, green: *cno* RNAi, red; *Dα-cat* RNAi, purple: *vinc* RNAi and light blue: *m6* RNAi). The correlation coefficient is 0.04. (h) Time-lapse images of Jub-GFP (upper) and myo-II-mKate2 (lower). Blue and magenta arrowheads indicate the decrease in Jub-GFP signal, and the formation of the PD junction, respectively. (i) Quantifications of Jub-GFP signal intensity around the rsMC based on time-lapse data captured at 25–26 h APF in *WT* cells. The Jub-GFP signal intensity along the line ROI at each position along the PD axis is plotted. Line colors represent the timing of measurement as shown in the upper left corner, where red and magenta lines correspond to 5 min before the rsMC formation and the time of rsMC formation, respectively. Steel-Dwass test: *WT* vs. *jub* RNAi, P < 0.001, *WT* vs. *cno* RNAi, P < 0.001, *WT* vs. *Dα-cat* RNAi, P < 0.001, *WT* vs. *vinc* RNAi, P > 0.5 (c), *WT* vs. *jub* RNAi, P < 0.001, *WT* vs. *cno* RNAi, P < 0.001, *WT* vs. *Dα-cat* RNAi, P < 0.001, *WT* vs. *vinc* RNAi, P > 0.5 (d). Welch’s *t*-test: the second time point vs. the last time point, P < 0.001 (i); this result indicated that the drop of Jub-GFP signal intensity at the time of rsMC formation was statistically significant. The number of rsMCs (c–e, g) and the number of junctions examined (i) are indicated. Data are presented as box plots (c, d) and as the mean ± s.d. (i). Scale bars: 2 μm (a, f, h).

To further explore the involvement of Jub, Cno, and Dα-cat in rsMC dynamics, we analyzed their distributions in *wild-type (WT*) cells. We found that the Jub signal was weakened specifically at the shrinking and reconnecting junctions (Fig. 2h, i, Supplementary Video 4). The Cno and Dα-cat signals decreased to a lesser extent during the junction exchange (Supplementary Fig. 5). These findings suggested that Jub, Cno, and Dα-cat prevented the detachment of cortical myo-II during the early phase of cell rearrangement, and that tuning the level of junctional Jub is important for the correct formation of the rsMC and subsequent completion of junction reconnection.

### M6 is responsible for the reduced level of Jub during junction exchange

Next, we investigated the mechanism responsible for the reduction of Jub levels during the junction exchange. We have shown that actin interacting protein 1 (AIP1) and cofilin retained Jub, Cno and Dα-cat at the shrinking AP junctions to prevent the precocious formation of the rsMC (Supplementary Fig. 6a, b; *flare (flr*) and *twinstar (tsr*) encode *Drosophila aip1* gene and *cofilin* gene, respectively) (ref. 16 and this study). Thus, AIP1 and cofilin may be eliminated from the AJ immediately before junction exchange, leading to the reduction of Jub levels. However, both AIP1 and cofilin were present along the reconnecting junctions; a strong AIP1 signal was detected inside the rsMC, and the cofilin signal was maintained after the junction shrinkage (Supplementary Fig. 6c–e). These data indicated that the rsMC formation is not accounted for by the decrease in AIP1 and/or cofilin levels at short junctions.

Alternatively, we noticed that a rsMC-like structure was occasionally formed around the vertex of the non-remodeling junctions (Fig. 3a). This observation led us to a hypothesis that TCJ proteins^8,9^ reduce junctional Jub levels to promote the rsMC formation. To test this hypothesis, we performed loss-of-function experiments on TCJ proteins. We found that RNAi of *m6*, which is known to accumulate at the tricellular SJ (tSJ) and lipid rafts^22–24^, increased the fraction of long-lasting rsMCs, without inducing the drastic malformation of the rsMC (Fig. 3b–e, Supplementary Fig. 2h–j, Supplementary Video 5). Jub was persistently retained inside such long-lasting rsMCs (Fig. 3f, g, Supplementary Video 6). Taken together, our data strongly suggest that the TCJ protein, M6, weakens the linkage between myosin and AJ by reducing the Jub level.

**Fig. 3.**
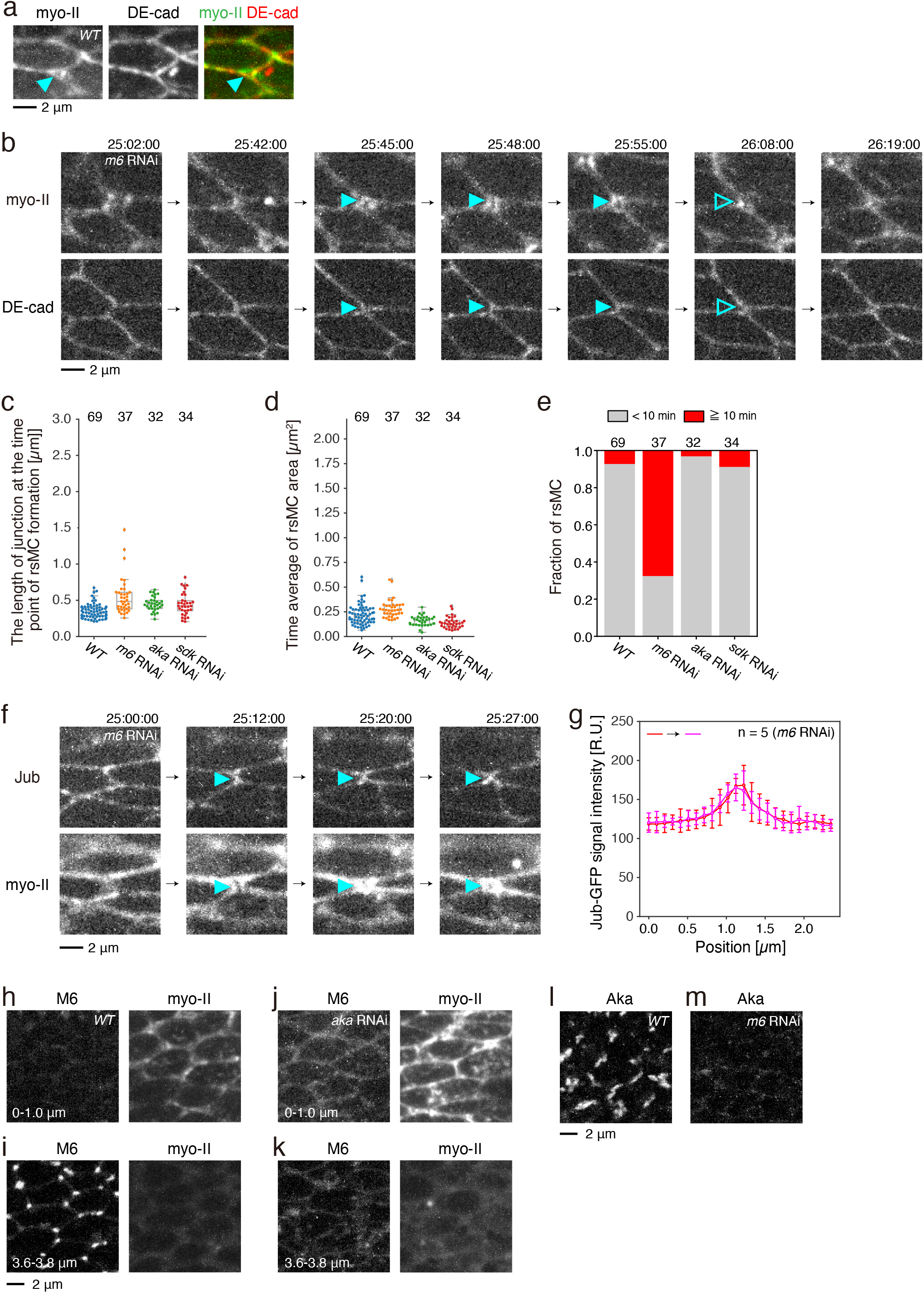
M6 is responsible for the reduced level of Jub during junction exchange. (a) Magnified images of myo-II-GFP (gray in left panel, green in right panel) and DE-cad-mTagRFP (gray in middle panel, red in right panel) at the tricellular junction (*i.e*., at the cell vertex) at 24 h APF. myo-II is occasionally detached from the vertex of non-remodeling junctions and forms a rsMC-like structure (arrowheads). (b) Time-lapse images of myo-II-mKate2 (upper) and DE-cad-GFP (lower) in *m6* RNAi wing. (c–e) Quantification of RNAi phenotypes. Junction length at the time point of rsMC formation (c), the temporal average of rsMC area (d), and the fraction of rsMC time duration (e) for each genotype based on time-lapse data captured at 25–26.5 h APF. (f) Time-lapse images of Jub-GFP (upper) and myo-II-mKate2 (lower) in *m6* RNAi wing. Arrowheads indicate the retention of Jub inside the rsMC. (g) Quantifications of Jub-GFP signal intensity around the rsMC based on time-lapse data captured at 25–26 h APF in *m6* RNAi cells (see the legend of Fig. 2i for details). (h–k) Images of M6-GFP (left panels) and sqh-mKate2 (right panels) at 24 h APF in *WT* (h, i) and *aka* RNAi (j, k) wings. Images along the AJ planes (h, j) and more basal planes (i, k) are shown as indicated at the bottom. (l, m) Images of Aka-GFP at 24 h APF in *WT* (l) and *m6* RNAi (m) wings. Steel-Dwass test: *WT* vs. *m6* RNAi, P < 0.001, *WT* vs. *aka* RNAi, P < 0.001, *WT* vs. *sdk* RNAi, P < 0.001 (c), *WT* vs. *m6* RNAi, P < 0.05, *WT* vs. *aka* RNAi, P > 0.05, *WT* vs. *sdk* RNAi, P < 0.01 (d). Welch’s *t*-test: *WT* vs. *m6* RNAi, P < 0.01; this result confirmed that the Jub-GFP signal intensity at the time point of rsMC formation was statistically different among *WT* and *m6* RNAi cells. The number of rsMCs (c–e) and the number of junctions examined (g) are indicated. Data are presented as box plots (c, d) and the mean ± s.d. (g). Scale bars: 2 μm (a, b, f, i, l).

It has been reported that M6 and the tSJ protein Anakonda (Aka) localize to the TCJ in a mutually dependent manner and that they cooperate in the TJC assembly in other epithelial tissues^23,25^. Thus, it is possible that Aka works with M6 in the junction exchange in the wing. However, we detected no strong phenotype of the rsMC in *aka* RNAi cells (Fig. 3c–e, Supplementary Fig. 2f, Supplementary Fig. 6l). The signal intensity of M6 was decreased upon *aka* RNAi, but M6 was still detected at vertices (Fig. 3h–k). In addition, M6 was shifted to the AJ plane, where it may act on Jub, in *aka* RNAi cells (Fig. 3j). On the contrary, the Aka signal was hardly detected in *m6* RNAi cells (Fig. 3l, m). The localization of Aka was more severely affected by *m6* RNAi, which might lead to the difference in the defect in the rsMC dynamics under the condition employed in this study.

RNAi of *sdk*, which encodes the tAJ protein Sdk^10–12^, did not strongly interfere with the rsMC dynamics (Fig. 3c–e, Supplementary Fig. 6m). We confirmed that the signal of Sdk-YFP dropped below the detection limit upon *sdk* RNAi (Supplementary Fig. 2g). The elongation of newly generated junctions was slowed down upon *sdk* RNAi in the wing (WT: 0.23 ± 0.20 μm/min, *sdk* RNAi: 0.09 ± 0.04 μm/min). This agrees with previous reports on male genitalia^10^. These results suggest that *sdk* RNAi is effective and that M6 and Sdk are responsible for different aspects of cell rearrangement.

### Interplay between M6 and Jub during junction exchange

Having established that M6 reduces the junctional Jub level during the junctional exchange, we investigated whether there is a counter-regulation from Jub to M6. M6 predominantly localized at the tSJ in *WT* (Fig. 4a–c). In *jub* RNAi cells, however, M6 was also detected at the AJ, indicating that M6 could stay at the AJ from which Jub was eliminated (Fig. 4d–f). Dlg remained localized in the plane basal to myo-II and that the Dlg signal was aligned along the apico-basal axis (Fig. 4g), suggesting that the basic structure of SJ was maintained in *jub* RNAi cells. M6 appeared at the AJ inside the enlarged rsMC of *flr* or *tsr* RNAi cells (Supplementary Fig. 6f–k), which is consistent with the loss of Jub along the remodeling junctions in these cells (Supplementary Fig. 6a, b). Collectively, our data uncovered an interplay between Jub and M6 during cell rearrangement.

**Fig. 4.**
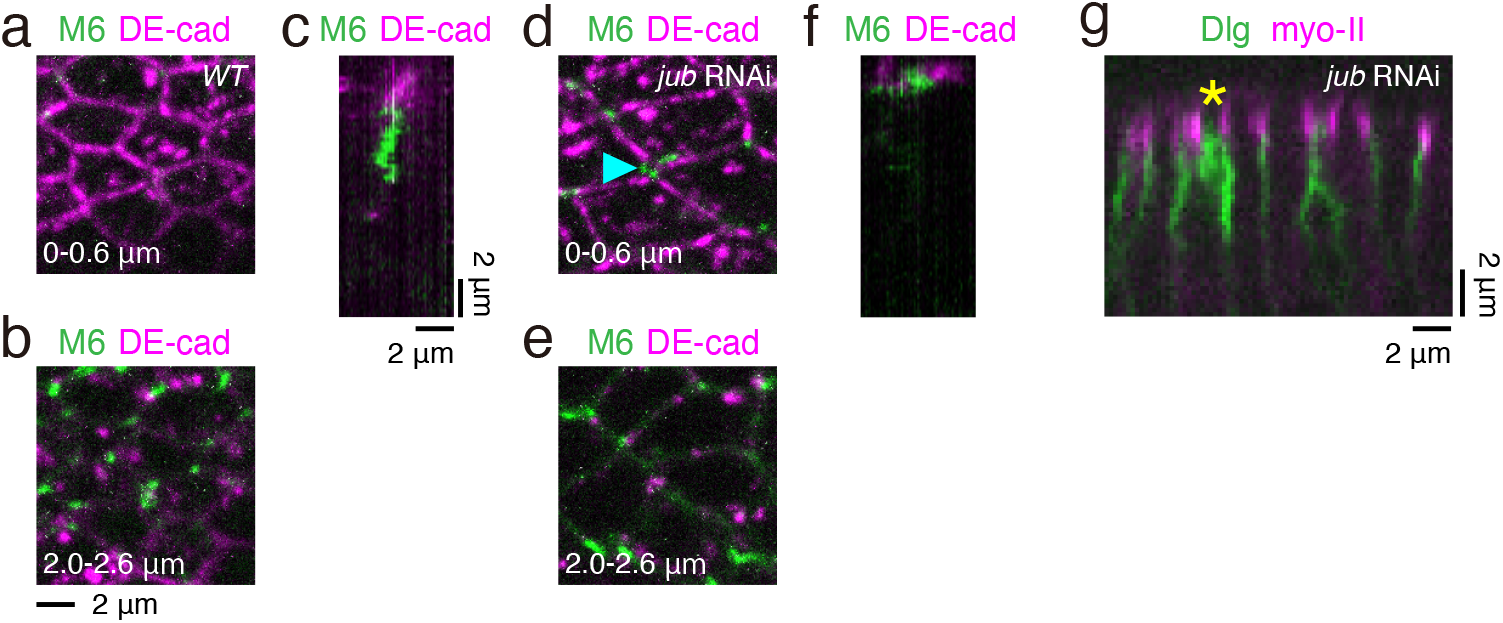
M6 is shifted to the AJ plane upon *jub* RNAi. (a–f) Images of M6-GFP (green) and DE-cad-mTagRFP (magenta) at 24 h APF in *WT* (a–c) and *jub* RNAi (d–f) wings. (a,b, d, e) Images along the AJ planes (a, d) and more basal planes (b, e) are shown as indicated at the bottom. (c, f) The side-view of short remodeling junctions. (g) Side-view of Dlg-GFP (green) and myo-II-mKate2 (magenta) at 24 h APF in *jub* RNAi wings. dsRNA against *jub* was expressed in the C region by using *ptc-Gal4* (Supplementary Fig. 2a). Asterisk indicates the boundary between the B region and C region. Scale bars: 2 μm (b, c, g).

Based on these findings, we propose that the junction shrinkage and elongation autonomously adjust the interplay between Jub and M6 to reconnect junctions. During the early phase of cell rearrangement, M6, which localizes at the vertex of the long remodeling junction, cannot exert a strong inhibitory effect on junctional Jub. Thus, Jub can maintain the linkage between cortical myo-II and the AJ. As cell rearrangement proceeds, the junction shrinkage brings the vertices at both ends of remodeling junction into proximity, which can elevate the inhibitory effect of M6 on Jub. Once the level of Jub starts to decrease, M6 gains more access to the AJ. This positive feedback leads to the detachment of cortical myo-II, specifically at short junctions. Subsequently, the elongation of the newly formed junction activates the feedback in the opposite direction, which allows the reattachment of cortical myo-II to junctions for the completion of junction reconnection.

### Junction geometry and mechanics govern the detachment of cortical myo-II cables for junction exchange

To understand the physical basis of junction exchange, we constructed a simple mechanical model based on the wetting theory^26^ and explored how changes in mechanical parameters, which are regulated by the molecular interactions described above, control the attachment and detachment of cortical myo-II cables to the junction. First, we considered the attached and detached states of cortical myo-II cables in a single cell (blue lines and red lines, respectively, in Fig. 5a). The myo-II cables are under tension and attached to the junction via the actin-AJ cross linkers. The free energy is originated from the cable’s tension and adhesion to the junction, whose energies per unit length are *γ* and –*σ*, respectively, and are uniform in the vicinity of the remodeling junction. Then, the energy cost of tension *γ* and energy gain of adhesion *σ* lead to the difference in the free energies of the attached and detached states: *F_d_* – *F_a_* = –2*l*(1 – cos *θ*)*γ* + *σ*(*L*_0_ + 2*l*) Fig. 5a, c–e for definitions and quantifications of geometrical quantities *L*_0_, *l*, and *θ*). The detached state is energetically preferable when *F_d_* is smaller than *F_a_*, which results in the rsMC formation.

**Fig. 5.**
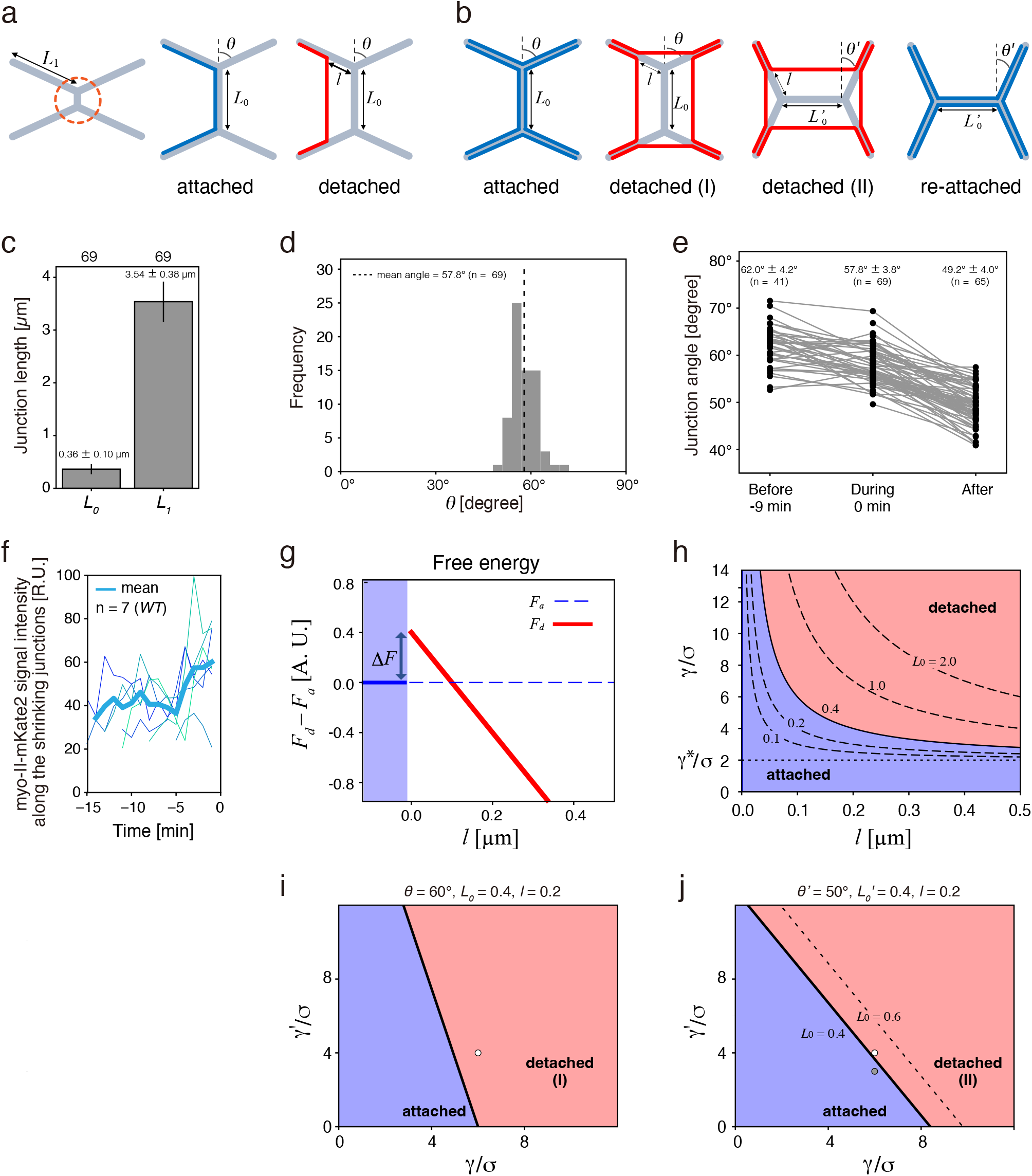
A mechanical model of the detachment/attachment of cortical myo-II. (a) Schematics of the attached and detached states of cortical myo-II cables in a single cell. An orange circle in the left panel indicates the region shown in the middle and right panels. In the attached state, myo-II cables, which are indicated by blue lines, run along cell contact surfaces. In the detached state, myo-II cables are detached from cell contact surfaces, as indicated by red lines. *L*_0_: the length of vertical remodeling junction; *L*_1_: the length of junctions connecting to the remodeling junction; *l*: the length of the connecting junctions, from which cortical myo-II is detached; *θ*: the half-angle between connecting junctions. (b) Schematics of the attached and detached states of cortical myo-II cables in four cells. 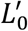 the length of horizontal newly generated junction; *θ′* + 90°: the angle between the connecting junction and the horizontal newly generated junction. (c–e) Quantification of the geometrical parameters of the theoretical model of myo-II cables. (c) Average length of the remodeling (*L*_0_) and connection junctions (*L*_1_) at the time point of rsMC formation in *WT* cells. For measuring *L*_1_, the average length of four connecting junctions was calculated for each rsMC. (d) Frequency of the half-angle between connecting junctions (*θ*) at the time point of rsMC formation. The average half-angle value of the upper and bottom connecting junctions was calculated for each rsMC. (e) Quantification of junction angles (*θ* and *θ′*) before, during, and after the rsMC formation (before: 9 min before the rsMC formation; during: the time point of the rsMC formation; after: 0–2 min after the rsMC fission). (f) Plot of myo-II-mKate2 signal intensity along shrinking AP junctions prior to the myo-II detachment. Bold line shows the average values. *t* = 0 indicates the time point when the myo-II gap was formed. (g) Free energies for the attached and detached states, *F_a_* and *F_d_*. Difference of the free energies *F_d_ – F_a_* is shown as a function of *l*. The energy scale is in arbitrary unit. The parameters are set as *γ* = 6.0, *σ* = 1.0, *θ* = 60°, and *L*_0_ = 0.4 *μm*. (h) A phase diagram of the attached state and detached state on a *γ/σ-l* plane. Solid and dotted lines indicate *γ = *σ*(L*_0_ + 2*l*)/2(1 – cos*θ*)*l* above which the detached state is energetically preferable. *γ*\* ≡ *σ*/(1 – cos *θ*) is the threshold tension for the detachment state to exist. *θ = 60°*. (i, j) Phase diagrams of the attached and detached states on the *γ/σ-γ′/σ* plane. Thick black lines indicate the conditions above which the detached state is preferable. The parameters are set as (i) *θ* = 60°, *L*_0_ = 0.4, and *l* =0.2, and (j) *θ′* = 50°, 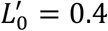, and *l* =0.2. In (j), the same condition for 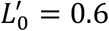 is also shown as a dotted line. Paired *t*-test: During vs. After, P < 0.001 (e). The number of rsMCs (c, d) and the number of junctions examined (e) are indicated. Data are presented as the mean ± s.d.

The analysis of the model indicated that an energy barrier Δ*F* = *σL*_0_ is present between the attached and detached states (Fig. 5g) and that the detached state is energetically preferable in the region *γ* > *σ*(*L*_0_ + 2*l*)/2(1 – cos *θ*)*l* with a threshold tension *γ*\* ≡ *σ*/(1 – cos *θ*) (Fig. 5h). Thus, large *γ* and/or small *σ* favor the detachment of myo-II cables from the junction. In addition, the conditions for the detachment depend on the geometrical quantities as well. For instance, the parameter region of the detached state is enlarged as *L*_0_ decreases (Fig. 5h). The half-angle between connecting junctions *θ* also controls the threshold tension *γ*\*; *γ*\* = *σ* when *θ* = 90° and *γ\** becomes infinitely large as *θ* reaches to zero. Taken together, the balance between tension *γ* and adhesion *σ*, in cooperation with the cell geometry, determines the attachment and detachment of myo-II cables.

The results of the model analysis provided an interpretation of the experimental data. First, the model prediction that the junction shrinkage (*i.e*., small *L*_0_) lowers the energy barrier and tension required for the detachment is consistent with the experimental observation that the detachment of myo-II cables is triggered specifically at short remodeling junctions. Second, the half-angle is maintained at approximately *θ ≈* 60° during junction shrinkage (the first and second rows of Fig. 5e), suggesting that the change in junction angle is not a main trigger of myo-II detachment in wing cells. Third, as cell rearrangement proceeded, myo-II, which can be regarded as a proxy for active stress^27^ contributing to cable tension *γ*, was enriched (Fig. 5f)^28^ and Jub, which contributed to the adhesion energy *σ*, was eliminated (Fig. 2i). Together, the balance between *γ* and *σ* shifted to favor the detached state. Fourth, the model predicts that when *σ* is uniformly lowered at the junctions, cells can overcome the energy barrier Δ*F* = *σL*_0_ at large *L*_0_, which is consistent with the loss-of-function phenotypes of actin-AJ linkers (Fig. 2a–e).

Next, we considered myo-II cables in four cells surrounding reconnecting vertices (Fig. 5b). We assumed that detached myo-II cables maintain a rectangular shape. The tension at the vertical (AP) and horizontal (PD) edges of the rectangular rsMC is set as *γ* and *γ′*, respectively. A similar calculation as described above leads to the condition required for the myo-II detachment as *γ*(1 – cos *θ*) + *γ′*(1 – sin *θ*) > *σ*(2 + *L*_0_/2*l*) for the detached state (I) in Fig. 5b (Methods). As discussed above, the transition to the detached state is induced by the decrease in *L*_0_ by the junction shrinkage and/or the increase in *γ/σ* (Fig. 5i). After the junctions are reconnected and the newly generated junction elongates, the condition of the myo-II detachment changes as 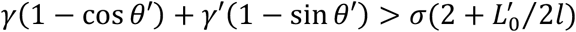 (the detached state (II) in Fig. 5b). According to this condition, the decrease in *γ* or *γ′* makes the attached state energetically favorable when the tension of the other type of edges of rsMC remains unchanged (Fig. 5j). Consistent with this model prediction, myo-II gradually decreased at the horizontal (PD) edges of myo-II cables, which eventually attached to the newly generated PD junction in *WT* cells (Fig. 1e).

Finally, we quantitatively compared our mechanical model with experimental data (see Methods for details). Based on the observation that the myo-II cable detachment occurred at *L*_0_ ≈ 0.4 μm in *WT* cells, we estimated (*γ,γ′*) ≈ (6*σ*, 4*σ*). The changes in the myo-II and Jub signal intensities at the shrinking AP junctions (Fig. 2i, Fig. 5f) suggest that the initial value of *γ* is below 3*σ*, with which the system is in the attached region at *θ* = 60° and *L*_0_ > 0.4 μm. Therefore, using the *WT* parameter values, cortical myo-II cables can be attached to the non-remodeling junction. After the short remodeling junctions were reconnected, *θ′* was approximately 50° at the time of rsMC fission (Fig. 5e). Substituting *θ′* = 50°, 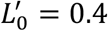 μm, and (*γ,γ′*) = (6*σ*, 4*σ*) into the condition equation for the detached state (II) indicates that the system resides at the boundary between the detached and attached states (Fig. 5j). Considering the decrease in the myo-II signal intensity along the horizontal edges of the rsMC (Fig. 1e) and the increase in 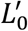 by the junction elongation, the parameter values are changed as (*γ,γ′*) ≈ (6*σ*, 3*σ*) and 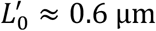. With the updated parameter values, a retransition from the detached state to the attached state is induced (Fig. 5j). These results suggest that the transition between the attached and detached states is possible using the estimated *in vivo* parameters.

In summary, the theoretical analysis of the minimal mechanical model elucidates how the physical properties of cortical myo-II cables are integrated with the junction geometry to govern their dewetting/wetting behaviors during cell rearrangement.

### Pten lowers tension of myo-II cables to induce its reattachment to the junction

To further clarify how temporal changes in tension affect the reattachment of myo-II cables to the junction, we analyzed *phosphatase and tensin homolog (pten*) RNAi cells. In the *Drosophila* wing, PTEN is responsible for lowering the levels of phosphatidylinositol (3,4,5)-triphosphate and myo-II at the PD junctions after junction exchange, leading to the elongation of newly generated PD junctions^28^. We found that myo-II levels were elevated and the reduction in myo-II levels at the horizontal (PD) edges of the rsMC was suppressed in *pten* RNAi cells (Fig. 6a). Under such conditions, the myo-II cables could not attach to the AJ (Fig. 6b–e, Supplementary Video 7). The comparison of the myo-II signal intensity between *WT* and *pten* RNAi cells yielded estimates of the ratio of cable’s tension and adhesion (*γ,γ′*) ≈(9*σ*, 7*σ*) (see Methods for details). Substituting the estimated parameter values into the condition equation for the detached state (II) confirmed that reattachment was prohibited in *pten* RNAi cells. These results further supported the validity of the proposed mechanical model.

**Fig. 6.**
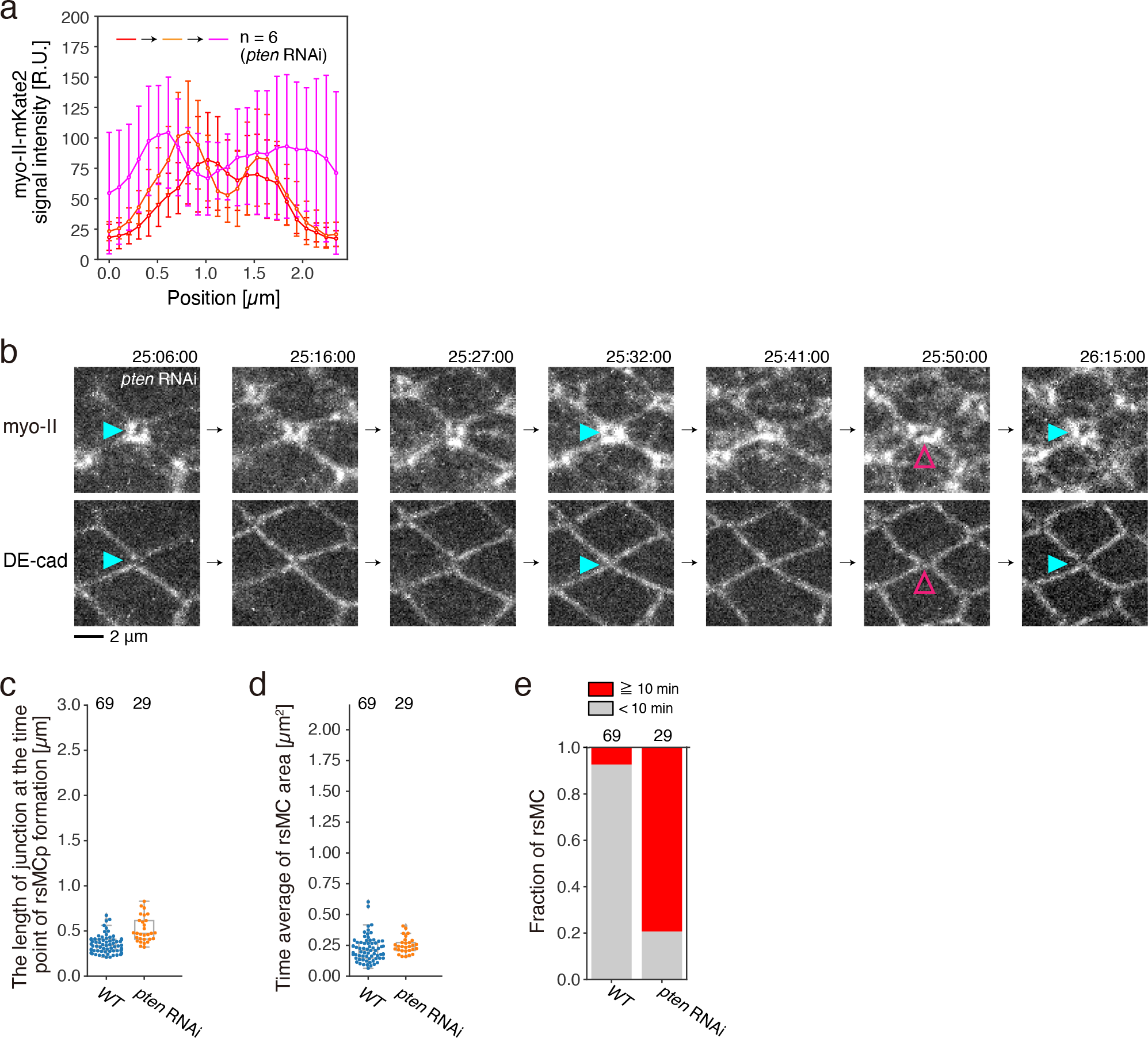
Cortical myo-II did not reattach to the AJ in *pten* RNAi cells. (a) Quantifications of myo-II-mKate2 signal intensity around the rsMC based on time-lapse data captured at 25–26.5 h APF at 25 °C in *pten* RNAi cells. The myo-II-mKate2 signal intensity along the line ROI, which aligns along the horizontal (PD) axis, is plotted (see the legend of Fig. 1e for details). (b) Time-lapse images of myo-II-mKate2 (upper) and DE-cad-GFP (lower) in *pten* RNAi wing. Blue and magenta arrowheads point to the rsMC. (c–e) Quantification of RNAi phenotypes. Junction length at the time point of rsMC formation (c), the temporal average of rsMC area (d), and the fraction of rsMC time duration (e) for each genotype based on time-lapse data captured at 25–26.5 h APF. Steel-Dwass test: *WT* vs. *pten* RNAi, P < 0.001 (c), *WT* vs. *pten* RNAi, P > 0.5 (d). The number of ROIs (a) and the number of rsMCs examined (c–e) are indicated. Data are presented as the mean ± s.d. (a) and box plots (c, d). Scale bar: 2 μm (b).

## Discussion

This study delineates the mechanism by which junctions are exchanged in epithelial cells (Fig. 7). The size and duration time of the rsMC are autonomously coordinated through the junction geometry and mechanics to loosen the junctional structure locally and transiently. The transient formation of the rsMC may prevent dilution of DE-cad at loosened junctions as suggested by the decrease of DE-cad signal at long-lasting rsMCs (magenta arrowheads in Fig. 2a). In addition, the directional information of the shrinking junction is inherited by the differential myo-II accumulation along the vertical and horizontal edges of the rsMC, which ensures unidirectional cell rearrangement by inducing the reattachment to junctions along edges with lower myo-II levels. This represents an advantage of the rsMC–mediated mechanism over *in situ* regulation at cell–cell contact surfaces in supporting the fidelity of junction exchange.

**Fig. 7.**
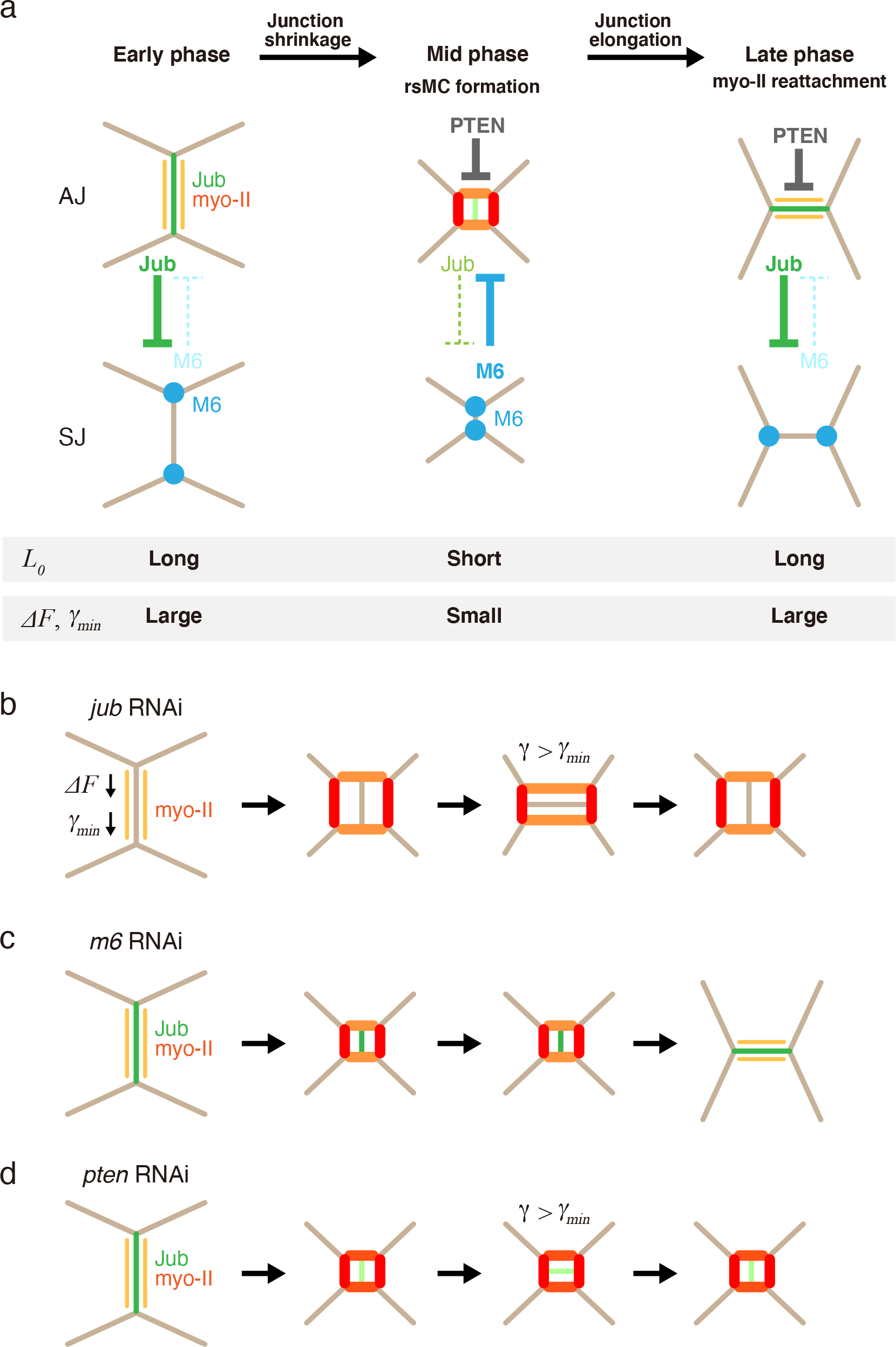
Summary and working hypothesis of the mechanism of junction exchange. (a) A schematic illustrating molecular interactions and the junction geometry during three phases in cell rearrangement. The concentrations of myo-II and Jub are depicted by color intensity (myo-II: light orange to red, Jub: light green to green). *L*_0_: the length of vertical remodeling junction; Δ*F*: an energy barrier for the detachment of cortical myo-II (= *σL*_0_); *γ_min_*: the value of tension of myo-II cables to separate the detached and attached regions (= *σ*(*L*_0_ + 2*l*)/2(1 – cos *θ*)*l*). (b–d) Schematic illustrating molecular interactions and the junction geometry in the indicated genotypes (b, *jub* RNAi; c, *m6* RNAi and d, *pten* RNAi). See the main text for details.

M6 is not detected in immature epithelial cells, including germband cells, and cells start to express M6 as the epithelium matures^12^. Thus, immature epithelial cells such as germband cells may deform cell membranes via actin-rich basolateral protrusions to form a multicellular rosette^7^. In contrast, mature epithelial cells such as pupal wing cells may reconnect junctions at more apical sides via M6. The mechanism whereby M6 regulates AJ components such as Jub in wing cells (this study) and Cno in Ras^V12^-overexpressing eye disc cells^22^ remains unclear. Since the mammalian homologue of M6 is involved in the actin regulation^24,29^, M6 may control actin remodeling, which is known to affect the structure and dynamics of AJ^16,19,30^. Alternatively, M6 may directly interact with the AJ components through a rapid, short-term translocation to the AJ (K.I. and K.S., preliminary data), as the position of junction compartments can dynamically change during the development^31^. More experiments such as simultaneous tracking of the AJ and SJ components^23,25^ will be necessary to elucidate how AJ and SJ interact and coordinate during junction exchange.

We introduced a simple theoretical model and uncovered the mechanical and geometrical conditions under which cortical myo-II cables attach to and detach from the junction. These conditions are compatible with the temporal changes in myo-II and jub levels in *WT* and RNAi cells. However, our model considers only the static force balance and thus cannot describe the entire process of junction exchange. To understand the dynamics of junction exchange, a more comprehensive model, including the turnover of actomyosin and junction components, needs to be developed. Recently, Nestor-Bergmann et al. proposed the Apposed-Cortex Adhesion Model, which investigates how the duration of cell-cell membrane linkage at the molecular scale affects cell rearrangement^32^. Extending our model to incorporate the concept of Apposed-Cortex Adhesion Model is an interesting direction for future research.

In conclusion, the present study shed light on the orchestration of geometry, mechanics and signaling around the cell vertex for reconnection of junctions. The proposed mechanism can potentially contribute to other aspects of morphogenesis. For instance, the rsMC-like structure at the vertex of a non-remodeling junction implies its relevance in a ratchet-like movement of the vertex along the junction^13^. In addition, because loss-of-function of M6 causes excess extrusion of cancer cells^22^, it is possible that M6 and actin-AJ linkers function to sense the risk of fracture at cell vertices, thereby preventing the formation of wounds in the epithelium. Given recent findings on the reorganization of the cytoskeleton and information sensing at the cell vertex^8,9,13,15,33,34^, further dissecting the complex interplay between actin-AJ regulators, TCJ proteins, and the geometry and mechanics at the vertices will clarify the origin of collective cell behaviors in epithelial development and plasticity.

## STAR Methods

### KEY RESOURCES TABLE

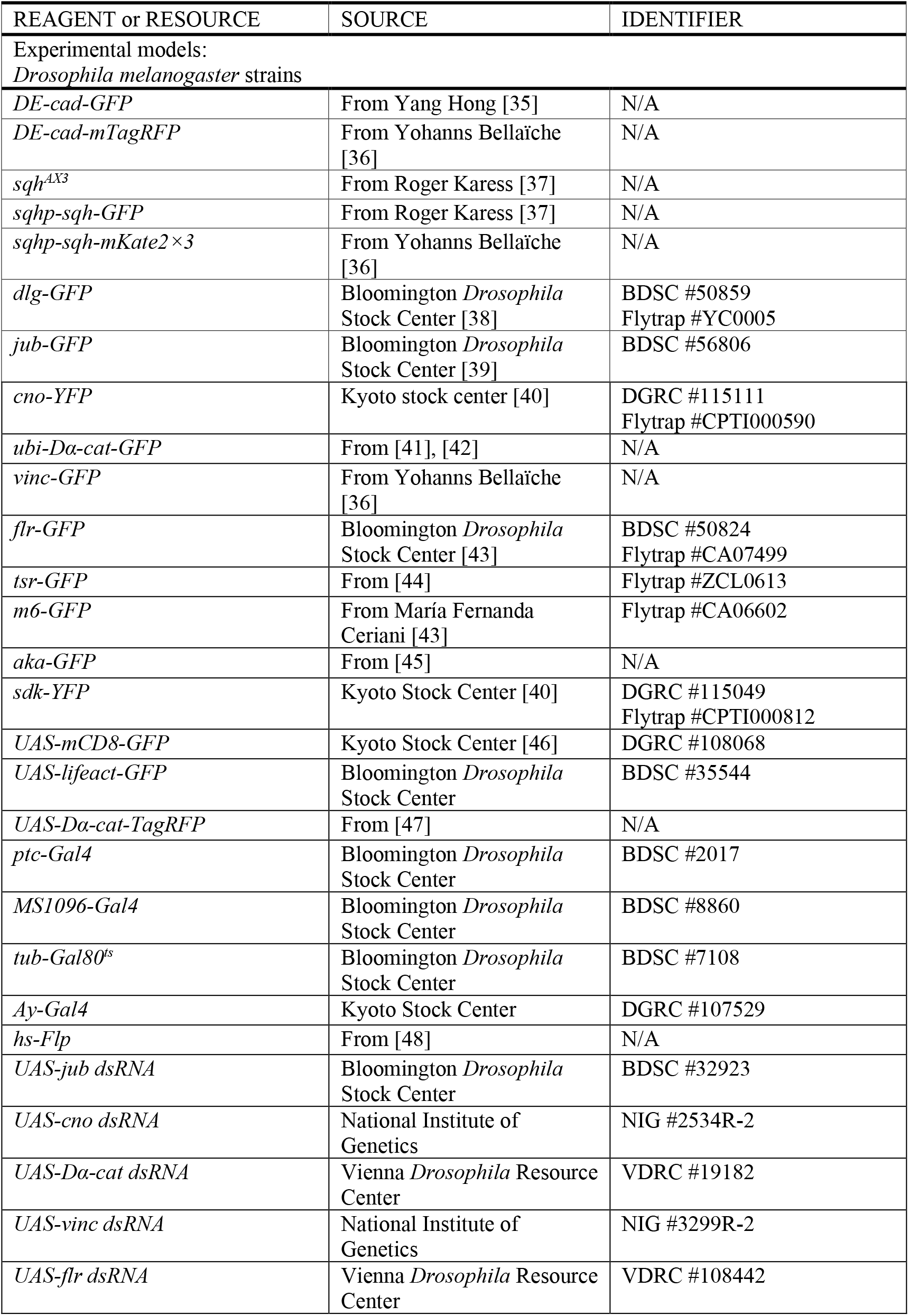
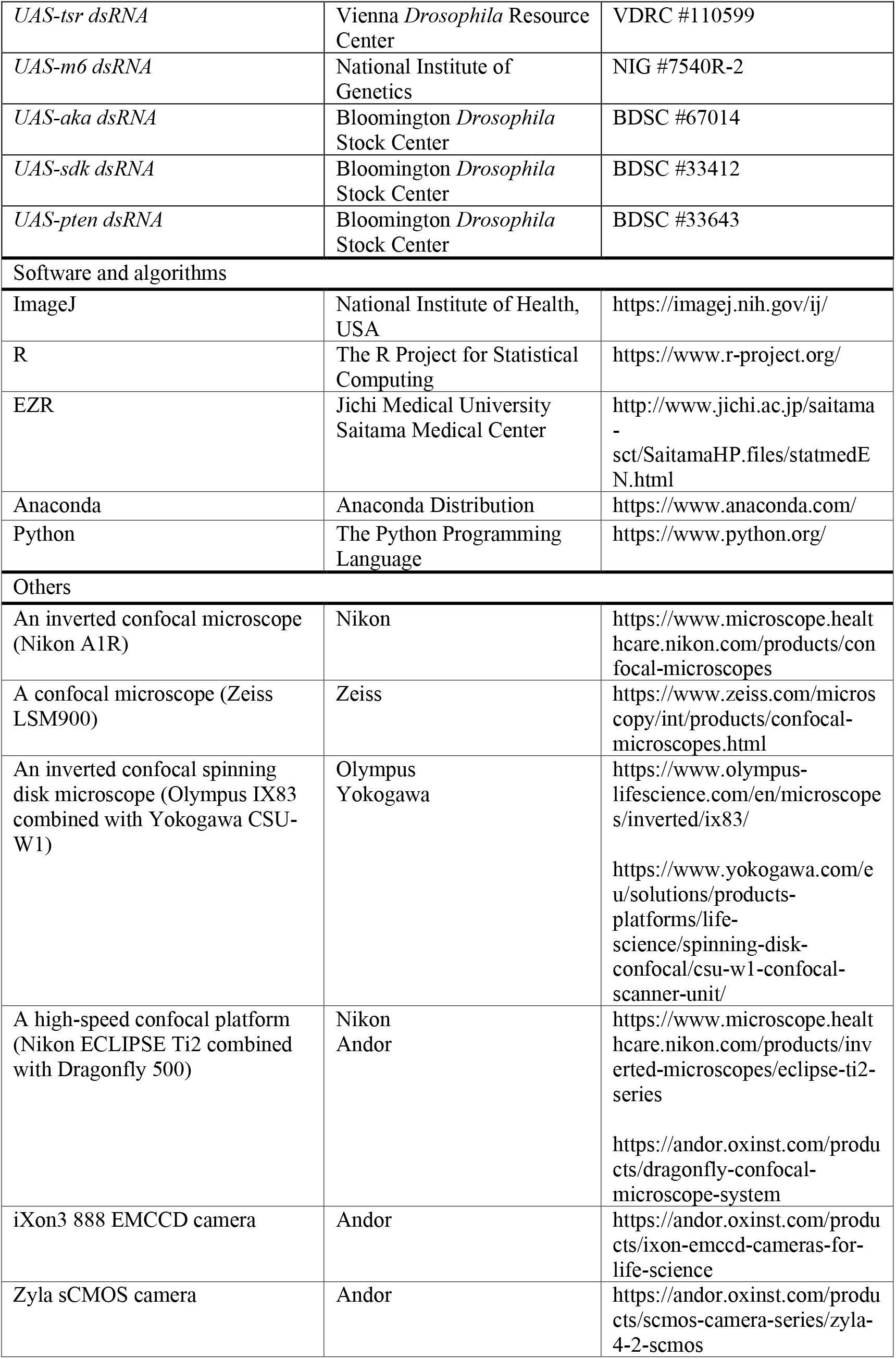

### RESOURCE AVAILABILITY

#### Lead contact

Further information and requests for resources and reagents should be directed to and will be fulfilled by the lead contact, Kaoru Sugimura.

#### Materials availability

This study did not generate new unique reagents.

#### Data and code availability

The authors declare that the data supporting the findings of this study are available within the paper and its Supplementary files. The data are available from the corresponding authors upon reasonable request.

### EXPERIMENTAL MODEL AND SUBJECT DETAILS

#### *Drosophila* genetics

*Drosophila* stocks were maintained at 25°C or 17°C. The flies used in the present study were listed in the KEY RESOURCES TABLE. The list of genotypes and crossing conditions are summarized in Genotypes of the experimental model.

To generate a mosaic clone expressing Lifeact-GFP, heat shock activated flip-out-Gal4-UAS system was used^48^. *hs-Flp; Ay-Gal4* females were crossed to *sqhp-sqh-mKate2×3, UAS-Lifeact-GFP* males at 25°C overnight. After the vial had been maintained for four days, heat-shock at 37°C was performed for 90 min. White pupae were then picked up and observed at 24 or 25 h APF.

#### Genotypes of the experimental model

Genotypes used for each figure are listed including temperature and hours after puparium formation.

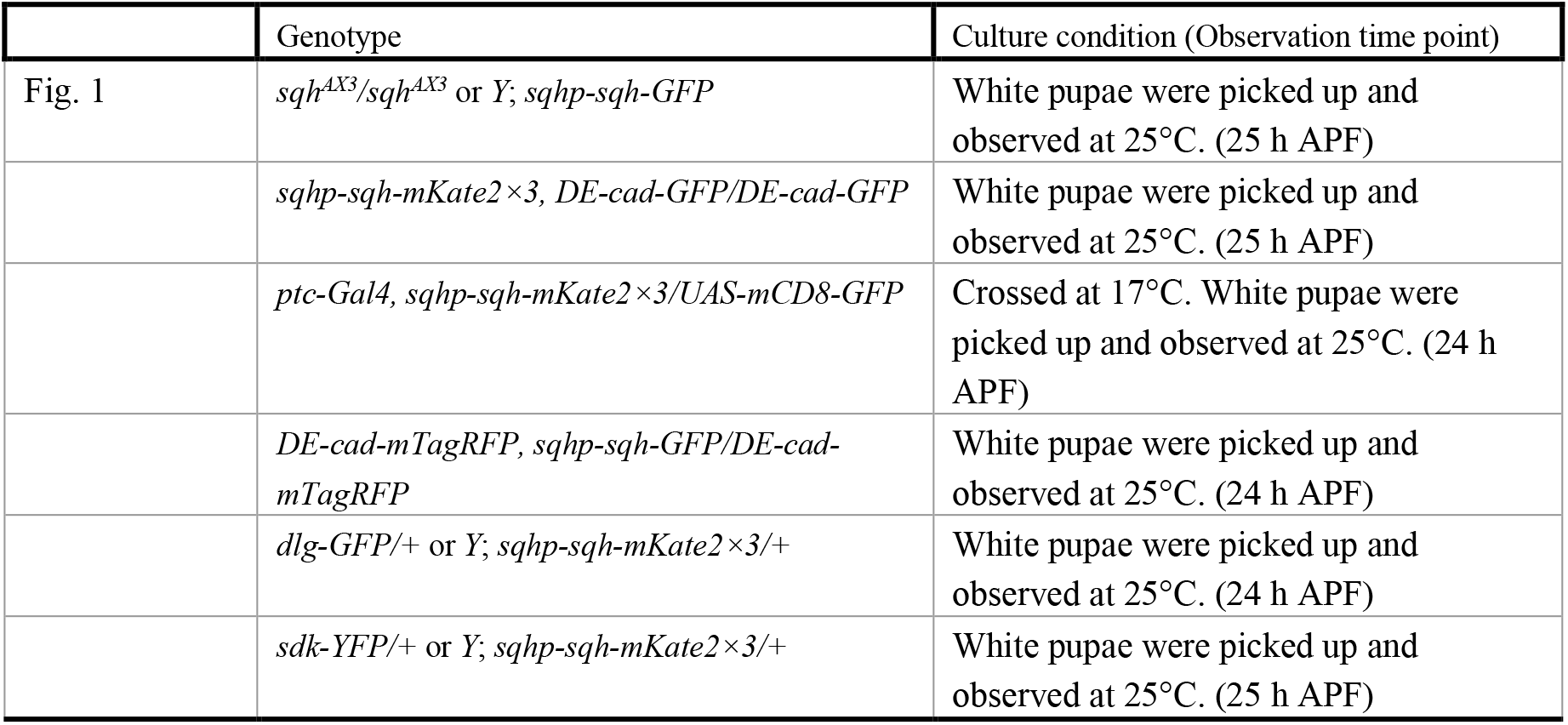
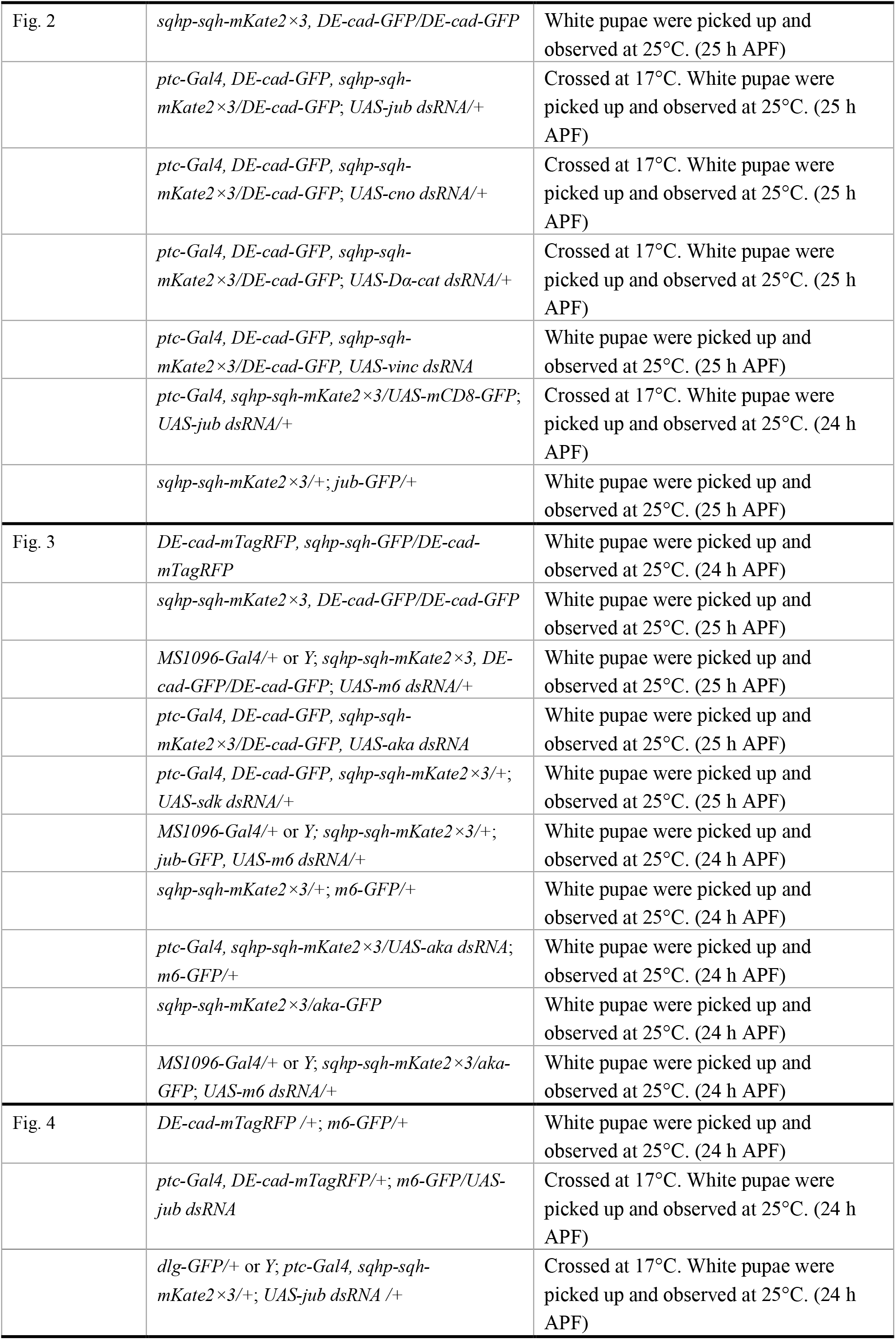
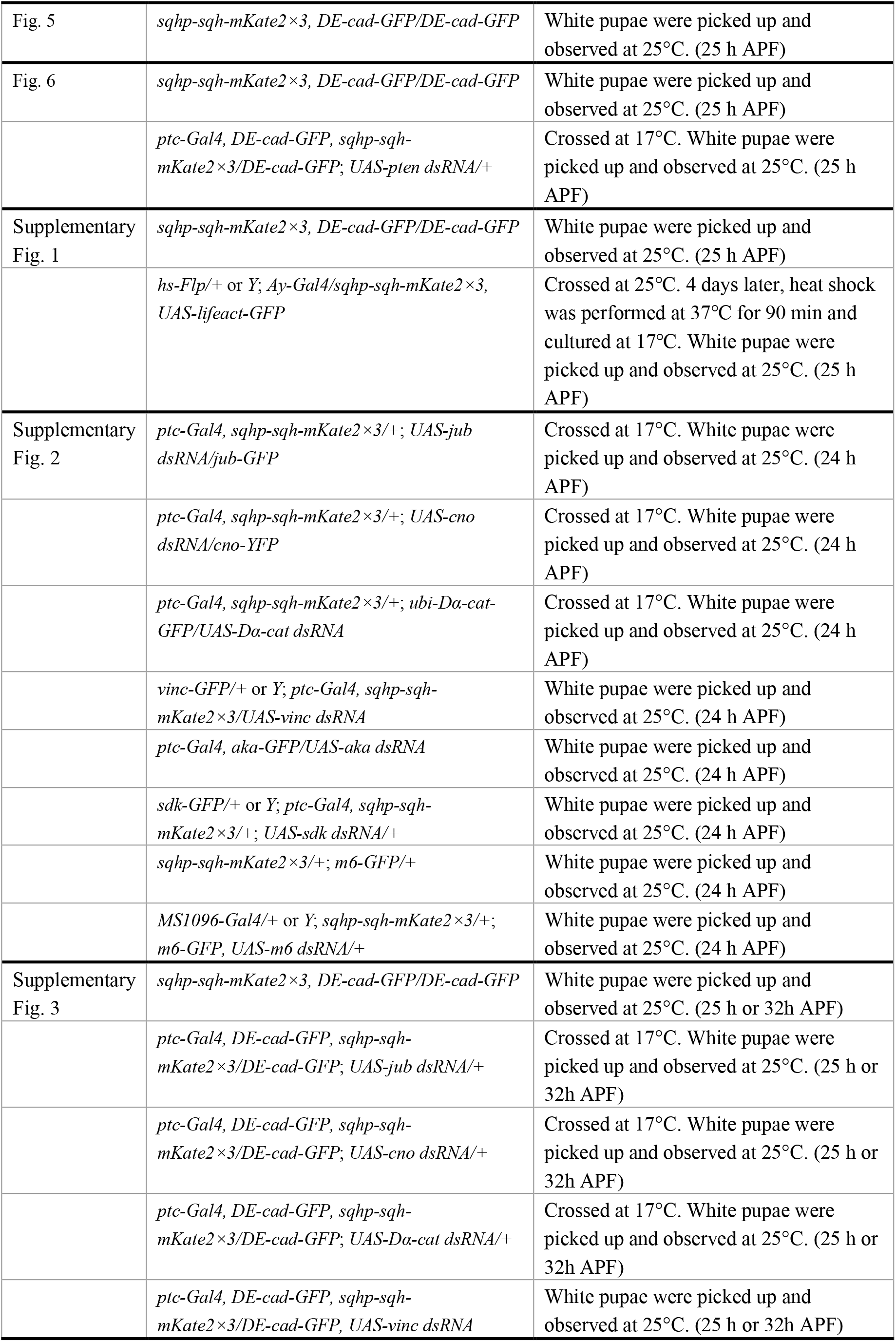
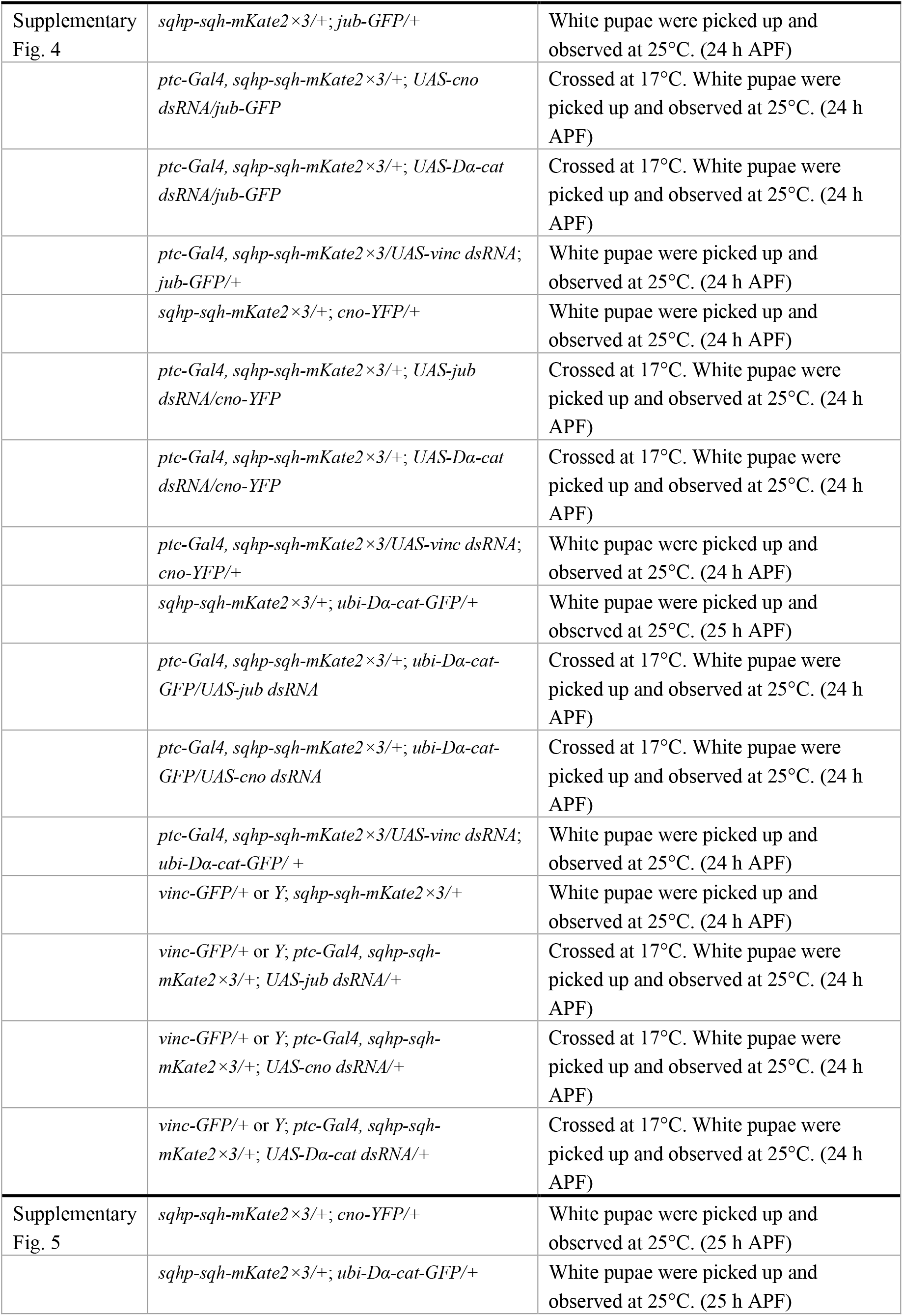
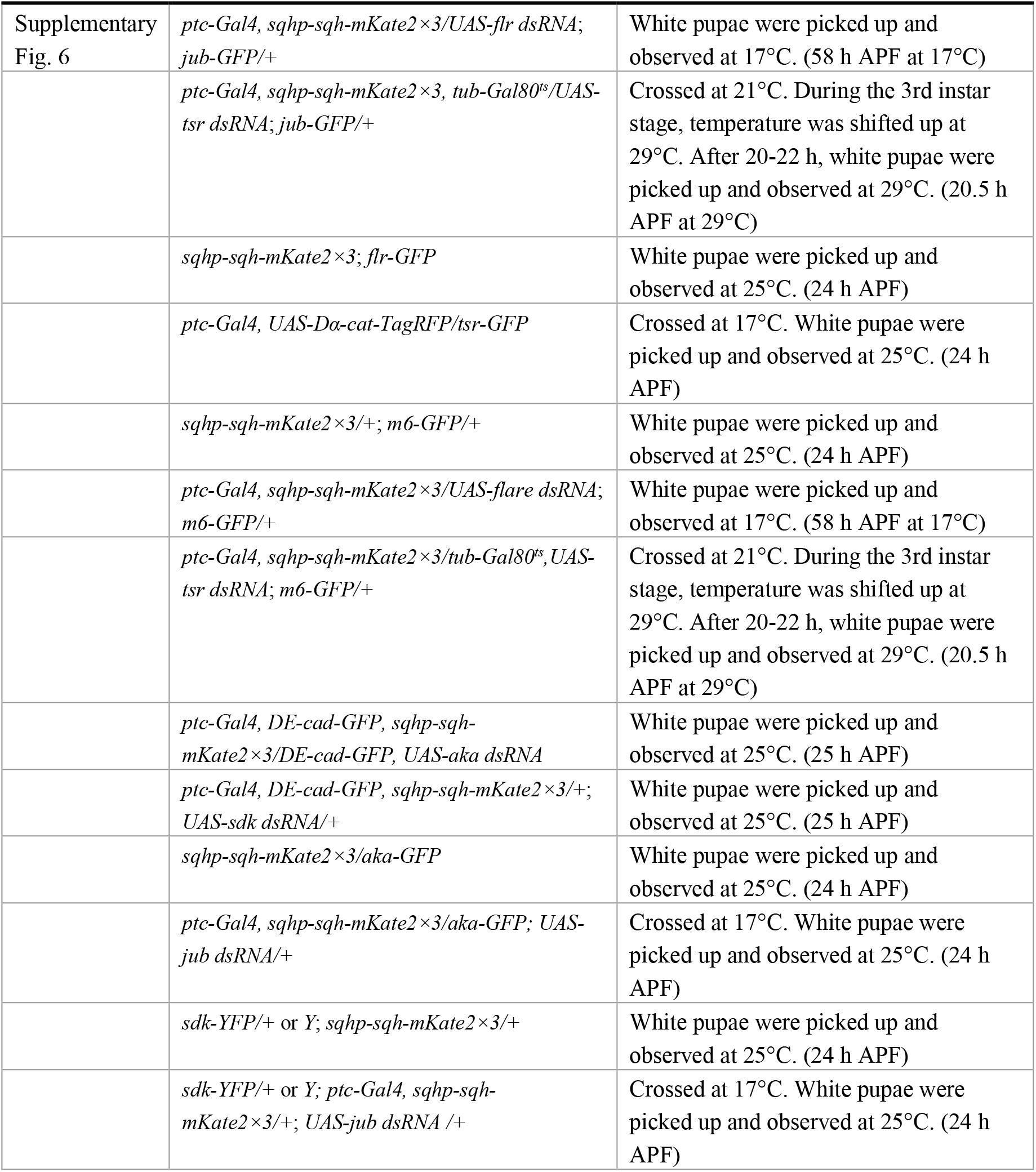

### METHOD DETAILS

#### Image collection

The preparation of *Drosophila* pupal wing samples for image collection was carried out as previously described^49,50^. In brief, pupae at appropriate ages were dissected to remove the pupal case on top of the left wing and were placed on a small drop of Immersol W 2010 (Zeiss 444969-0000-000) in a glass bottom dish. The fixed time-point images and cross section images shown in Fig. 1g–k, Supplementary Fig. 2b–g, i, j, and Supplementary Fig. 6e were acquired using an inverted confocal microscope (A1R; Nikon) equipped with a 60×/NA1.2 Plan Apochromat water-immersion objective at room temperature. Low-magnification images of the wing shown in Supplementary Fig. 3d–h were acquired using a confocal microscope (LSM 900; Zeiss) equipped with and a 10×/NA0.45 Plan Apochromat objective (Zeiss) at room temperature. The fixed time-point images shown in Fig. 4g and the time-lapse images shown in Supplementary Fig. 6c were acquired at 25°C using an inverted confocal spinning disk microscope (Olympus IX83 combined with Yokogawa CSU-W1) equipped with an iXon3 888 EMCCD camera (Andor), an Olympus 100×/NA1.35 SplanApo silicone oil-immersion objective, and a temperature control chamber (TOKAI HIT), using IQ 2.9.1 (Andor)^50^. Images in other figures were acquired using a high-speed confocal platform, Dragonfly (Nikon ECLIPSE Ti2 combined with Dragonfly 500 unit) equipped with an Zyla sCMOS camera (Andor), a 60×/NA1.20 CFI Plan Apochromat VC water-immersion objective (Nikon), using Fusion (Andor). After imaging, we confirmed that the pupae survived to at least the pharate stage.

#### Surgical manipulation to relax the tissue stretch

To analyze the effects of tissue tension on cell rearrangement (Supplementary Fig. 1f–h), wings were detached from the hinges using forceps. Details were described previously^16,49^. Briefly, before removing pupal case, hinges were severed at 23.5–24 h APF by rubbing forceps on hinge region. Pupal cases were then removed, pupae were mounted on glass bottom dishes as described image collection section.

### QUANTIFICATION AND STATISTICAL ANALYSIS

#### Image analysis

We focused on the rsMC generated from the AP junctions but not from the PD junctions. To quantify the area of the rsMC, the inner perimeter of the rsMC was manually tracked using the region of interest (ROI) manager in ImageJ. The junction length and the half-angle between connecting junctions were measured using line and angle tools, respectively, in ImageJ.

Z-projection was performed to extract the fluorescent signals on the AJ or SJ planes. The background signal of myo-II-mKate2 or DE-cad-GFP was subtracted using the ‘subtract background’ command (*r* = 50; except Supplementary Fig. 6d, see below), and the signal intensity was measured by manually selecting ROIs and using the ROI manager in ImageJ. The rsMC was divided into four edges (*v_1_, v_2_, h_1_*, and *h_2_*), as shown in Fig. 1d. In Fig. 1e and Fig. 6a, the spatial myo-II-mKate2 signal intensities at the rsMC were calculated using ‘Plot Profile’ in ImageJ with a line width of 10 pixel. In Fig. 2i and Fig. 3g, to measure the signal intensities of Jub-GFP, the background signal was subtracted using the ‘subtract background’ command (*r* = 100), and the signal intensity was quantified using ‘Plot Profile’ in ImageJ with a line width of 5 pixel. In Supplementary Fig. 6d, to measure the signal intensities of AIP1-GFP and myo-II-mKate2, the background signal was subtracted using the ‘subtract background’ command (*r* = 100), and the signal intensity was quantified using ‘Plot Profile’ in ImageJ with a line width of 7 pixel.

#### Statistics

P-values were calculated in R or Python based on the Mann-Whitney *U* test (Fig. 1d), the Steel-Dwass test (Fig. 2c, d, Fig. 3c, d, Fig. 6c, d, Supplementary Fig. 1c, d, f, g, Supplementary Fig. 3a, b), the Welch’s *t*-test (Fig. 2i, Fig. 3g), the correlation analysis (Fig. 2g, Supplementary Fig. 1i, j), and the paired *t*-test (Fig. 5e). The Steel-Dwass test was performed when comparing every pair in all groups. The Mann-Whitney *U* test, the Steel-Dwass test, and the Welch’s *t*-test were performed with EZR (Saitama Medical Center, Jichi Medical University, Saitama, Japan, https://www.jichi.ac.jp/saitama-sct/SaitamaHP.files/statmed.html), which is a graphical user interface for R (The R Foundation for Statistical Computing, Vienna, Austria)^51^. More precisely, it is a modified version of R commander designed to add statistical functions frequently used in biostatistics. The correlation analysis and paired *t*-test were performed with SciPy module in Python.

### MODELING

#### Mechanical model of the junction exchange

##### Model construction and analysis

We constructed a mechanical model of the cortical myo-II cables immediately before, during, and after the junction exchange. We considered the free energy, which consists of the tensile energy of the myo-II cables and the adhesive energy between the myo-II cables and junction. The underlying assumption is that the relaxation of tension and adhesion between myosin cables and AJ components is faster than the cell rearrangement time (~1 h) and rsMC dynamics (~5 min). Given that the lifetime of the binding of actin–AJ linkers to actin cytoskeleton to of the order of 1–10 s^52^, this assumption is valid for the situation considered here.

We assumed that detached myo-II cables maintain a rectangular shape and set up a symmetrical geometry for the sake of simplicity (Fig. 5a, b). Quantification of the junction geometry showed that the length of vertical remodeling junction *L*_0_ is approximately 0.4 μm (Fig. 5c). *l* is the length of the connecting junctions, from which cortical myo-II is detached. Since *l* was too small to measure precisely, we used an approximate value *l* ≈ 0.2 μm. Due to limitations in image resolution, the myo-II signals along the vertical remodeling junction and at the 0.2 μm-end of connection junctions could not be separated. We thus assume that the tension of myo-II cables does not change at the 0.2 μm-end of connection junctions.

The free energies of the attached and detached states (*F_a_* and *F_d_*) are originated from the cable’s tension and adhesion to the junction, whose energies per unit length are *γ* and –*σ*, respectively. The detachment of cortical myo-II cables is conditioned by *F_d_* – *F_a_* < 0. Before the vertical, AP junction is reconnected, the difference in the free energies of attached and detached (I) states are *F_d_* – *F_a_* = –4*γl*(1 – cos *θ*) – 4*γ′l*(1 – sin *θ*) + 2*σ*(*L*_0_ + 4*l*). This leads to the condition equation for the detachment *γ*(1 – cos *θ*) + *γ′*(1 – sin *θ*) > *σ*(2 + *L*_0_/2*l*) (Fig. 5i). After the reconnection of junctions, the reattachment is favored when 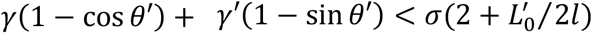 is satisfied, where *γ′* is the tension of horizontal edges of the rsMC (Fig. 5j).

##### Evaluation of model parameters

Following the data presented in Fig. 5c, we set *L*_0_ = 0.4 in Fig. 5g–j. The half-angle between the connecting junctions (*θ*) was approximately 60° at the time of the rsMC formation (Fig. 5d). Quantification of the change in *θ* during junction shrinkage (Fig. 5e) suggests that the change in junction angle is not the main trigger of myo-II detachment. After junction reconnection, as the newly generated junction elongates, *θ′* decreases, as shown in Fig. 5e. Based on these results, we treated *θ* and *θ′* as constants in the main text: *θ* = 60° for Fig. 5g–i, and *θ′* = 50° for Fig. 5j.

Experimental data showed that the ratio of myo-II-mKate2 signal intensity was ~3:2 for the vertical and horizontal edges of rsMC at the time of rsMC formation (Fig. 1d) and that the myo-II cable detachment occurred at *L*_0_ ≈ 0.4 μm (Fig. 2c). By substituting *γ* = (3/2)*γ′* and *in vivo* geometrical parameters into the condition equation for the detached state (I) in Fig. 5b, we estimated (*γ, γ′*) ≈ (6*σ*, 4*σ*) at the time point of rsMC formation (white circle in Fig. 5i). We use this value as a reference for parameter estimation. During the shrinkage of AP junctions, the myo-II signal intensity approximately doubled and the Jub signal intensity decreased by approximately 20% (Fig. 2i, Fig. 5f). Thus, we estimate that *γ* is initially below 3*σ*, with which the system is in the attached region at *θ* = 60° and *L_0_ >* 0.4 (Fig. 5h, i). During the junction exchange, the myo-II signal intensity along the horizontal edges of the rsMC dropped by approximately 20%, whereas that along the vertical edges remained unchanged (Fig. 1e). This observation provides the estimates of (*γ,γ′*) ≈ (6*σ*, 3*σ*) (gray circle in Fig. 5j). In addition, the length of the newly generated PD junction, 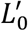 is increased during cell rearrangement. With (*γ, γ′*) ≈ (6*σ*, 3*σ*) and 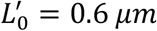, the re-attached state is favored energetically (Fig. 5j). We estimated model parameters in RNAi cells in the same procedure. Based on the myo-II signal intensity (magenta line in Fig. 6a), we estimated 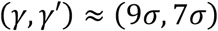 in *pten* RNAi cells. With these parameters, the reattachment of myo-II cables is prohibited even at *L*_0_ = 0.6 (Fig. 5j).

## Acknowledgments

The authors would like to thank Yohanns Bellaïche, Yang Hong, Roger Kares, María Fernanda Ceriani, the Bloomington Stock Center, the Kyoto Stock Center, the Vienna *Drosophila* Resource Center (VDRC), NIG-Fly, the *Drosophila* Genomics Resource Center, and the Developmental Studies Hybridoma Bank for reagents; Goshi Ogita, Miho Aruga, Kyoko Komano, Risa Matsui, and Mayu Miyakawa for technical assistance; the iCeMS Analysis Center, Shizue Ohsawa, and the Nagoya University Live Imaging Center for imaging equipment; and Tetsuhisa Otani and Yohanns Bellaïche for critical reading of the manuscript. This study was financially supported by JSPS KAKENHI Grant (17K15125), the AMED PRIME program (20gm5810025h9904), the Sumitomo Foundation (200303), and the Takeda Science Foundation to K.S., JSPS KAKENHI Grant (19K16139) and The Uehara Memorial Foundation (202110172) to K.I., JST CREST (JPMJCR1923) to S.I., and JSPS KAKENHI Grant (18KK0234) to Y.T.

## Author contributions

K.I. and K.S. designed the research. K.I. performed the experiments. Y.T. assisted the experiments. K.I., K.S and S.I. analyzed the data. S.I. constructed and analyzed mathematical model with input from K.S. K.S., K.I. and S.I. drafted the manuscript. All authors approved the final manuscript.

## Supplemental Information

**Supplementary Fig. 1.**
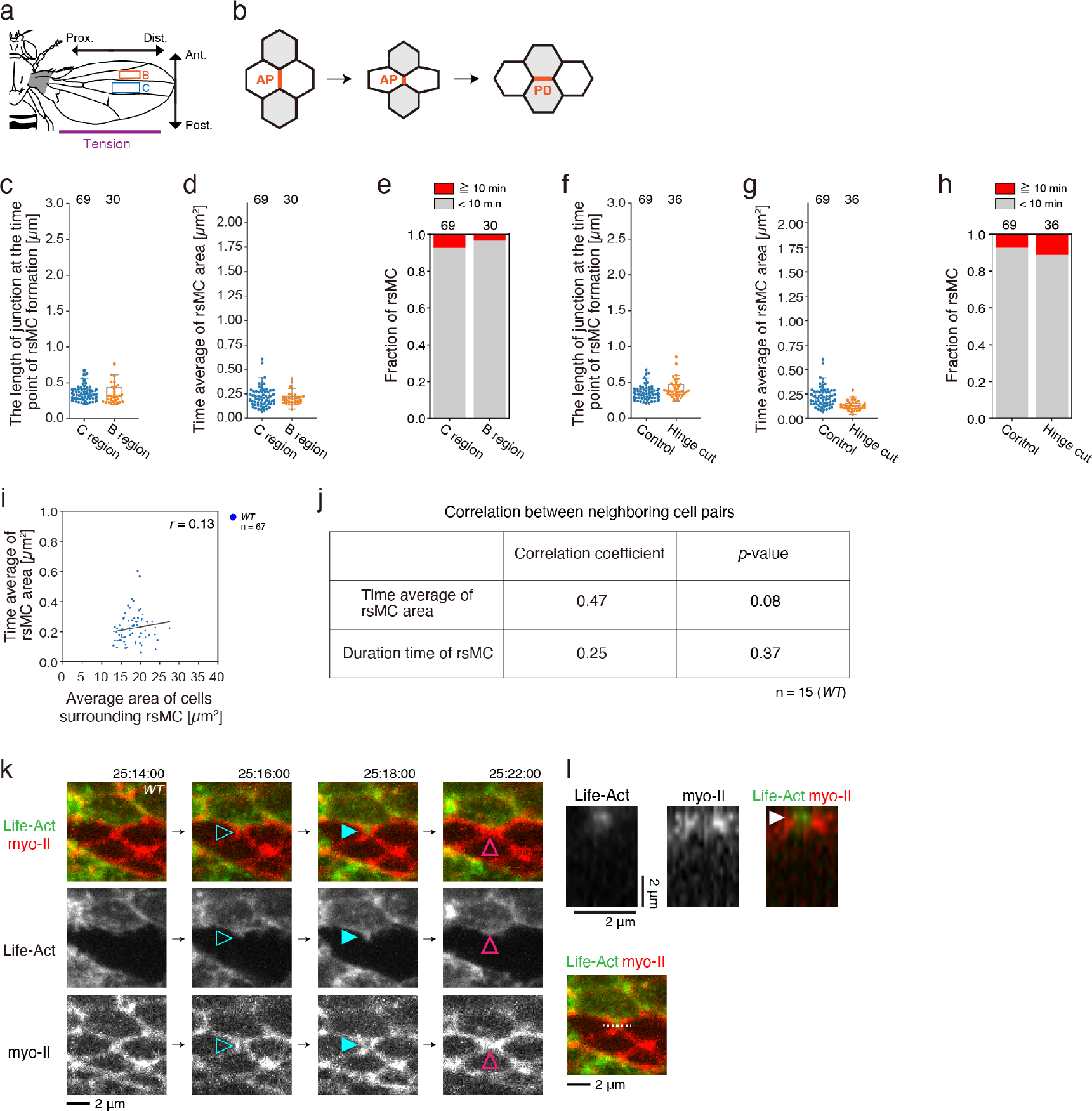
Characterization of rsMC morphology and dynamics. We analyzed the dependence of rsMC morphology and dynamics on different aspects of wing cells. First, by analyzing the movie data presented in Fig. 2c–e, we quantified the morphology and dynamics of rsMC in the B region of the wing. This analysis showed that rsMC behaviors were not influenced by its position in the tissue. Second, in the absence of extrinsic stretching force from the hinge, the rsMC was formed from a junction of similar length, but the rsMC was slightly smaller compared with the control. This suggests that rsMC formation *per se* does not depend on the extrinsic stretching force. We speculate that once myo-II cables are detached from the AJ, the extrinsic stretching force might enlarge the rsMC until the intrinsic and extrinsic forces are balanced. Third, the time average of rsMC area did not depend on the average area of cells surrounding the rsMC. Fourth, when the rsMC was formed sequentially in neighboring cells, rsMC morphology and dynamics were not strongly correlated among cell pairs. Overall, our data suggest that rsMC formation is locally regulated at every event, and that the tissue-level, extrinsic effect is small. It has been shown that actin-rich basolateral protrusions play pivotal roles in cell rearrangement during the germband elongation in *Drosophila* embryos^7^. We addressed whether this is also true in the *Drosophila* wing. Prior to the rsMC fission and the formation of a new junction, actin protrusions invaded the rsMC at the AJ, whereas no significant actin-rich protrusions were detected at the basolateral side of the cells. (a) Schematics of the adult fly (adapted from ref. 16). The hinge is shaded gray. In this and all subsequent figures, the vertical and horizontal directions are aligned with the AP and PD axes, respectively. We studied the intervein C region indicated by blue rectangle unless otherwise noted. We analyzed wings at 24–27 h APF when the PD cell rearrangement proceeds and cell division hardly occurs. Purple lines indicate the orientation of tissue tension generated by the hinge constriction^49,53^. (b) Wing cells rearrange along the PD axis at 21 h APF and afterward. AP junction shrinks and is remodeled to form a new PD junction. Orange lines indicate remodeling junctions. (c–e) Quantification of rsMC morphology and dynamics in the B and C regions of the *WT* wing (orange and blue rectangles in (a), respectively) based on time-lapse data captured at 25–26.5 h APF. Junction length at the time point of rsMC formation (c), the temporal average of rsMC area (d), and the fraction of rsMC time duration (e) are shown. (f–h) Quantification of rsMC morphology and dynamics in the C region of the control and mechanically relaxed wing based on time-lapse data captured at 25–26.5 h APF. Junction length at the time point of rsMC formation (f), the temporal average of rsMC area (g), and the fraction of rsMC time duration (h) are shown. (i) The time average of the rsMC area was plotted against the average area of cells surrounding each rsMC. The correlation coefficient is shown at the right upper corner. (j) Table of correlation coefficients for the temporal average of rsMC area and the duration time of rsMC among neighboring cell pairs. (k) Time-lapse images of Lifeact-GFP (green in upper panels, gray in middle panels) and myo-II-mKate2 (red in upper panels, gray in bottom panels). Lifeact-GFP was clonally expressed. (l) Side-view of Lifeact-GFP (gray in upper left panel, green in upper right panel) and myo-II-mKate2 (gray in upper middle panel, red in upper right panel) along the dashed line in bottom panel. Steel-Dwass test: C region vs. B region, P > 0.5 (c), C region vs. B region, P > 0.5 (d), Control vs. Hinge cut, P > 0.1 (f), Control vs. Hinge cut, P < 0.001 (g). The number of rsMCs (c–i) and the number of neighboring cell pairs examined (j) are indicated. Data are presented as box plots (c, d, f, g). Scale bars: 2 μm (k, l).

**Supplementary Fig. 2.**
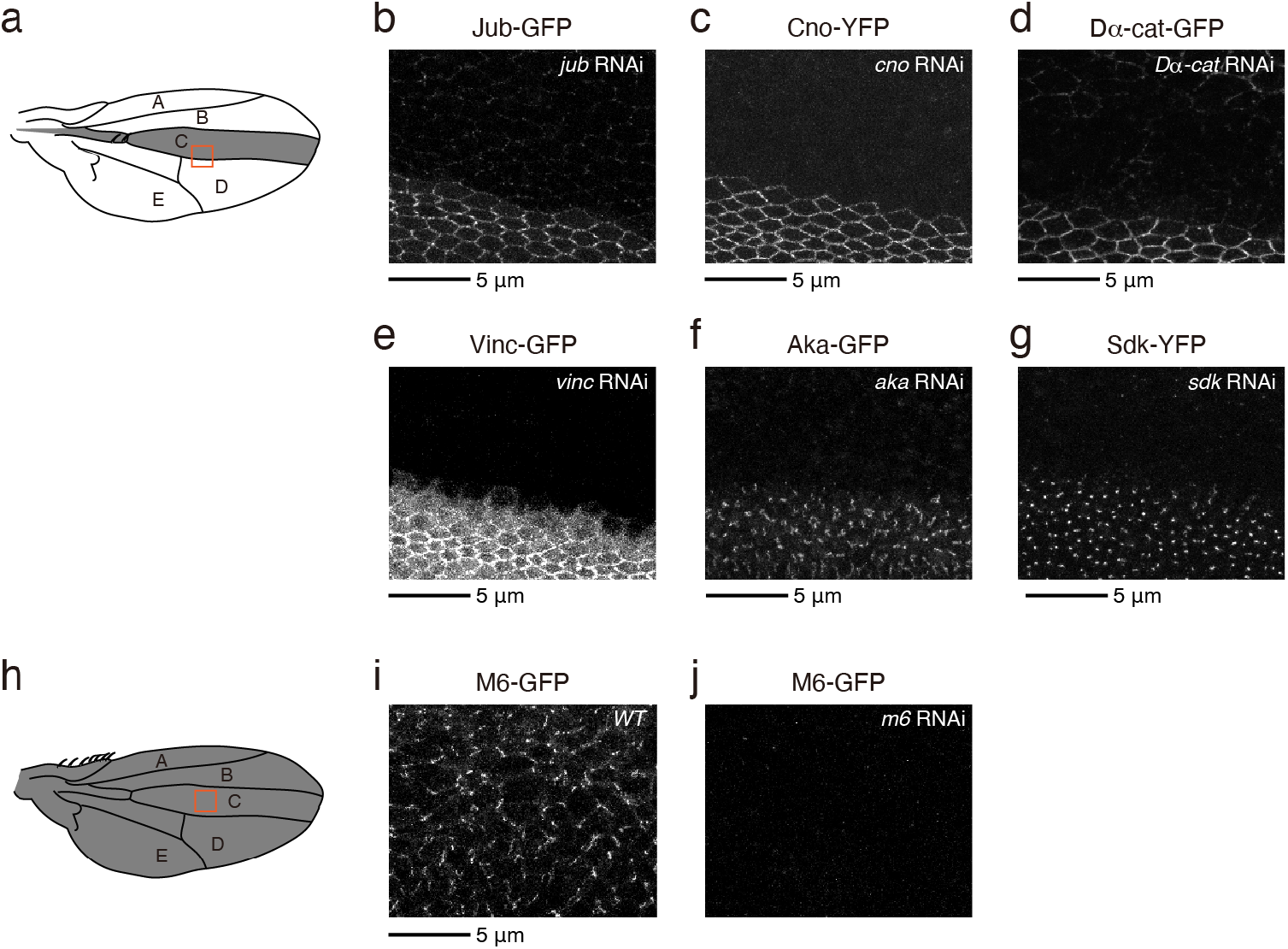
Characterization of RNAi lines used in this study. (a) Schematic of the wing. The expression domain of *ptc-Gal4* was shaded gray. Orange rectangle indicates the region shown in (b–g). (b–g) Images of Jub-GFP (b), Cno-YFP (c), Dα-cat-GFP (d), Vinc-GFP (e), Aka-GFP (f), or Sdk-GFP (g) at 24 h APF in wings of the indicated genotypes (b, *jub* RNAi; c, *cno* RNAi; d, *Dα-cat* RNAi; e, *vinc* RNAi; f, *aka* RNAi and g, *sdk* RNAi). dsRNA against each gene was expressed under the control of *ptc-Gal4*. (h) Schematic of the wing. The expression domain of *MS1096-Gal4* was shaded gray. Orange rectangle indicates the region shown in (i, j). (i, j) Images of M6-GFP at 24 h APF in wings of the indicated genotypes (i, *WT* and j, *m6* RNAi). In (j), dsRNA against *m6* was expressed under the control of *MS1096-Gal4*. We tested *MS1096-Gal4* and *ptc-Gal4* for expressing dsRNA against *m6*. We presented the data obtained by using *MS1096-Gal4*, which decreased the signal intensity of M6-GFP under the detection limit and yielded stronger phenotypes. *vinc* RNAi, *aka* RNAi or *sdk* RNAi did not induce the malformation of rsMC by using either of *MS1096-Gal4* or *ptc-Gal4*. Scale bars: 5 μm (b–g, i).

**Supplementary Fig. 3.**
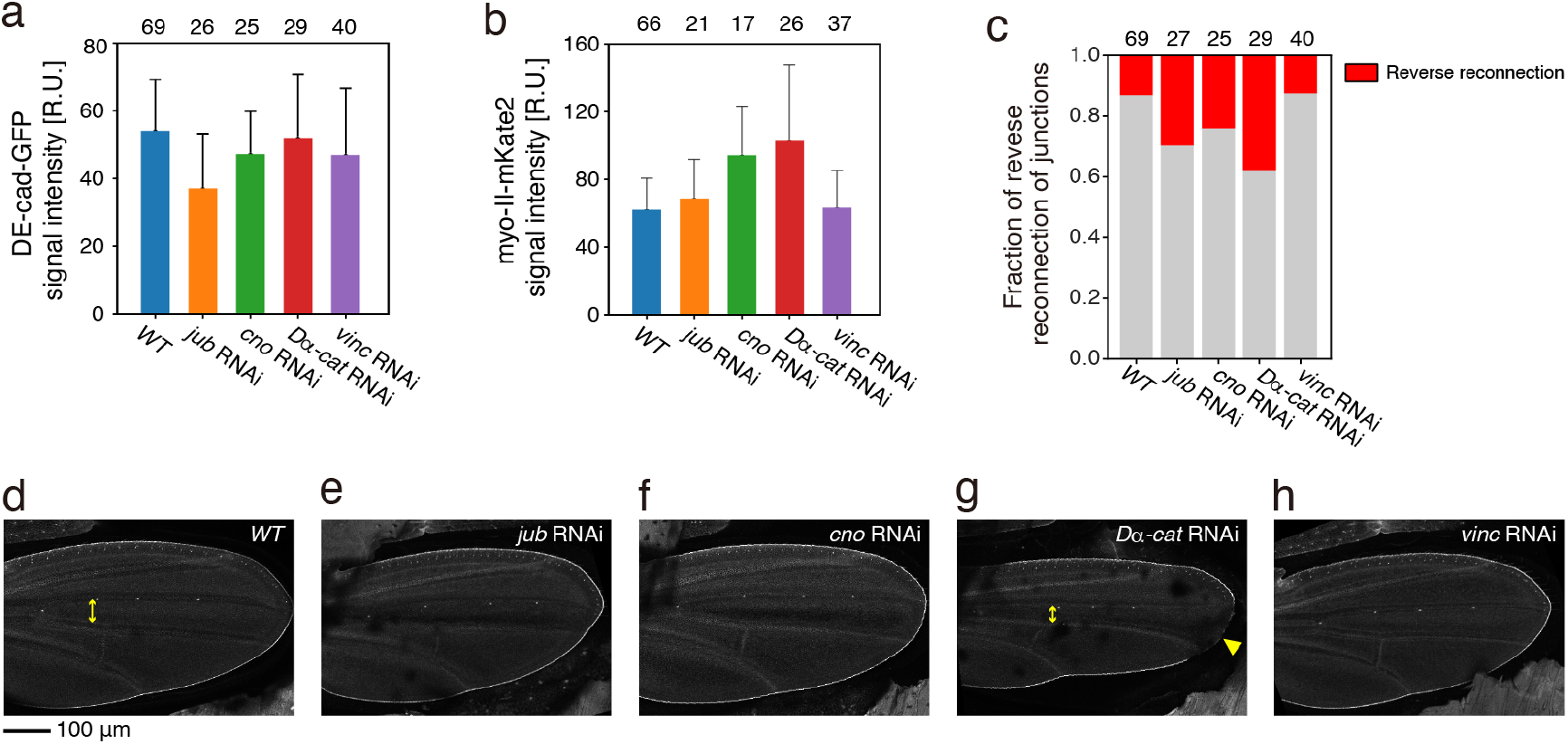
RNAi phenotypes of actin-AJ linkers. We examined phenotypes induced by RNAi of actin-AJ linkers. First, the signal intensity of DE-cad and myo-II hardly changed upon *vinc* RNAi, whereas *jub* RNAi decreased the DE-cad level and *cno* and *Dα-cat* RNAi increased the myo-II level. *jub* RNAi also resulted in the increase in the gap in junctional DE-cad at 27 h APF (WT: 5.5/698 cells and *jub* RNAi: 28/352 cells). These results may imply that the role of Jub is different from that of Cno and Dα-cat during the junction exchange, though other possibilities cannot be excluded. Second, RNAi of *jub*, *cno*, or *Dα-cat* resulted in the reverse reconnection of junctions in 30–40% of cases. Third, the whole-wing shape was not severely affected by *jub* or *cno* RNAi. Based our experience in the study of wing development, the defect in cell rearrangement is often difficult to detect at the whole-wing scale, when dsRNA was expressed only in the C region by using *ptc-Gal4*. In addition, the penetrance of the reverse reconnection of junctions may explain why the defect in the whole-wing shape was not detected. The C region was narrower along the AP axis in *Dα-cat* RNAi cells. We had seen a *Dα-cat* RNAi-like wing when the cell number in the C region was decreased by the defect in cell division. (a) The DE-cad-GFP signal intensity along the remodeling AP junctions at the time point of the rsMC formation for indicated genotypes. (b) The myo-II-mkate2 signal intensity along the remodeling AP junctions at 3 min before the rsMC formation for indicated genotypes. (c) Fraction of reverse reconnection of junctions. Reverse reconnection is defined as reconnection of the AP short remodeling junction inside the rsMC to form a new PD junction, then reconnecting again to form a new AP junction (see Fig. 2a). (d–h) Low magnification images of DE-cad-GFP of the *WT* (d), *jub* RNAi (e), *cno* RNAi (f), *Dα-cat* RNAi (g), and *vinc* RNAi (h) wings at 32 h APF. (g) The C region in the *Dα-cat* wing was narrower along the AP axis than in the control wing (double-pointed arrows). Developed nick in the wing’s distal margin (arrowhead). Steel-Dwass test: *WT* vs. *jub* RNAi, P < 0.001, *WT* vs. *cno* RNAi, P > 0.1, *WT* vs. *Dα-cat* RNAi, P > 0.5, *WT* vs. *vinc* RNAi, P < 0.05 (a), *WT* vs. *jub* RNAi, P > 0.5, *WT* vs. *cno* RNAi, P < 0.001, *WT* vs. *Dα-cat* RNAi, P < 0.001, *WT* vs. *vinc* RNAi, P > 0.5 (b). The number of junctions (a, b) and the number of rsMCs (c) examined are indicated. Data are presented as the mean ± s.d. (a, b). Scale bar: 100 μm (d).

**Supplementary Fig. 4.**
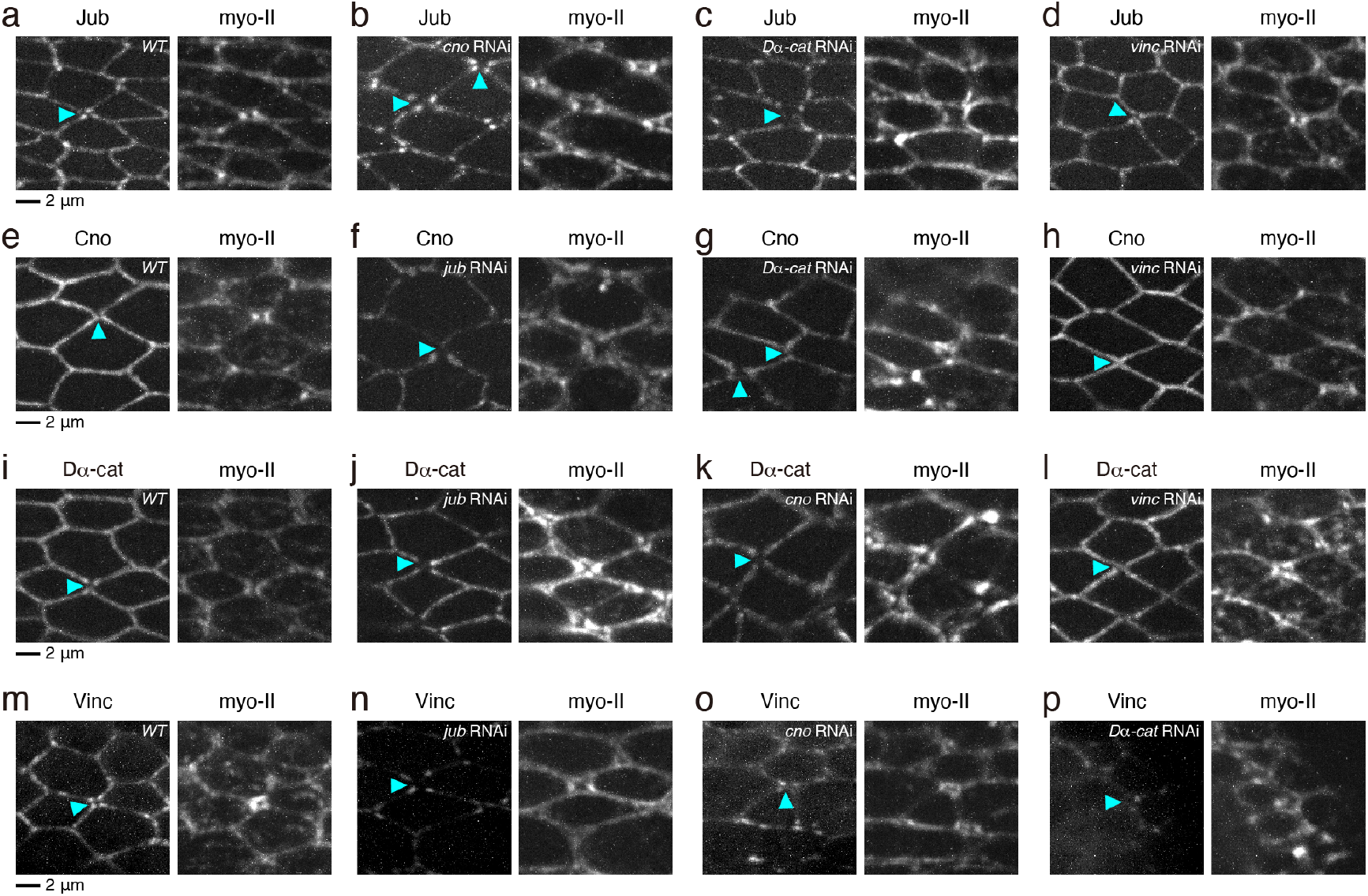
The dependence of actin-AJ linkers in their localization. To address why *vinc* RNAi did not significantly interfere with the rsMC formation and reattachment of cortical myo-II to junctions, we examined the distributions of Jub, Cno, and Dα-cat in *vinc* RNAi cells. We found that the distributions of Jub, Cno, and Dα-cat were largely normal in *vinc* RNAi cells. In contrast, Vinc was lost in *jub, cno, Dα-cat* RNAi cells. Thus, these data suggest that Vinc is in the downstream of Jub, Cno, and Dα-cat, and that Vinc is dispensable for the junction recruitment and activity of Jub, Cno, and Dα-cat. (a–d) Images of Jub-GFP (left) and myo-II-mKate2 (right) in *WT* (a), *cno* RNAi (b), *Dα-cat* RNAi (c), *vinc* RNAi (d) wings at 24 h APF. (e–h) Images of Cno-YFP (left) and myo-II-mKate2 (right) in *WT* (e), *jub* RNAi (f), *Dα-cat* RNAi (g), *vinc* RNAi (h) wings at 24 h APF. (i–l) Images of Dα-cat-GFP (left) and myo-II-mKate2 (right) in *WT* (i), *jub* RNAi (j), *cno* RNAi (k), *vinc* RNAi (l) wings at 24 h APF. (m–p) Images of Vinc-GFP (left) and myo-II-mKate2 (right) in *WT* (m), *jub* RNAi (n), *cno* RNAi (o), *Dα-cat* RNAi (p) wings at 24 h APF. Scale bars: 5 μm (a, e, i, m).

**Supplementary Fig. 5.**
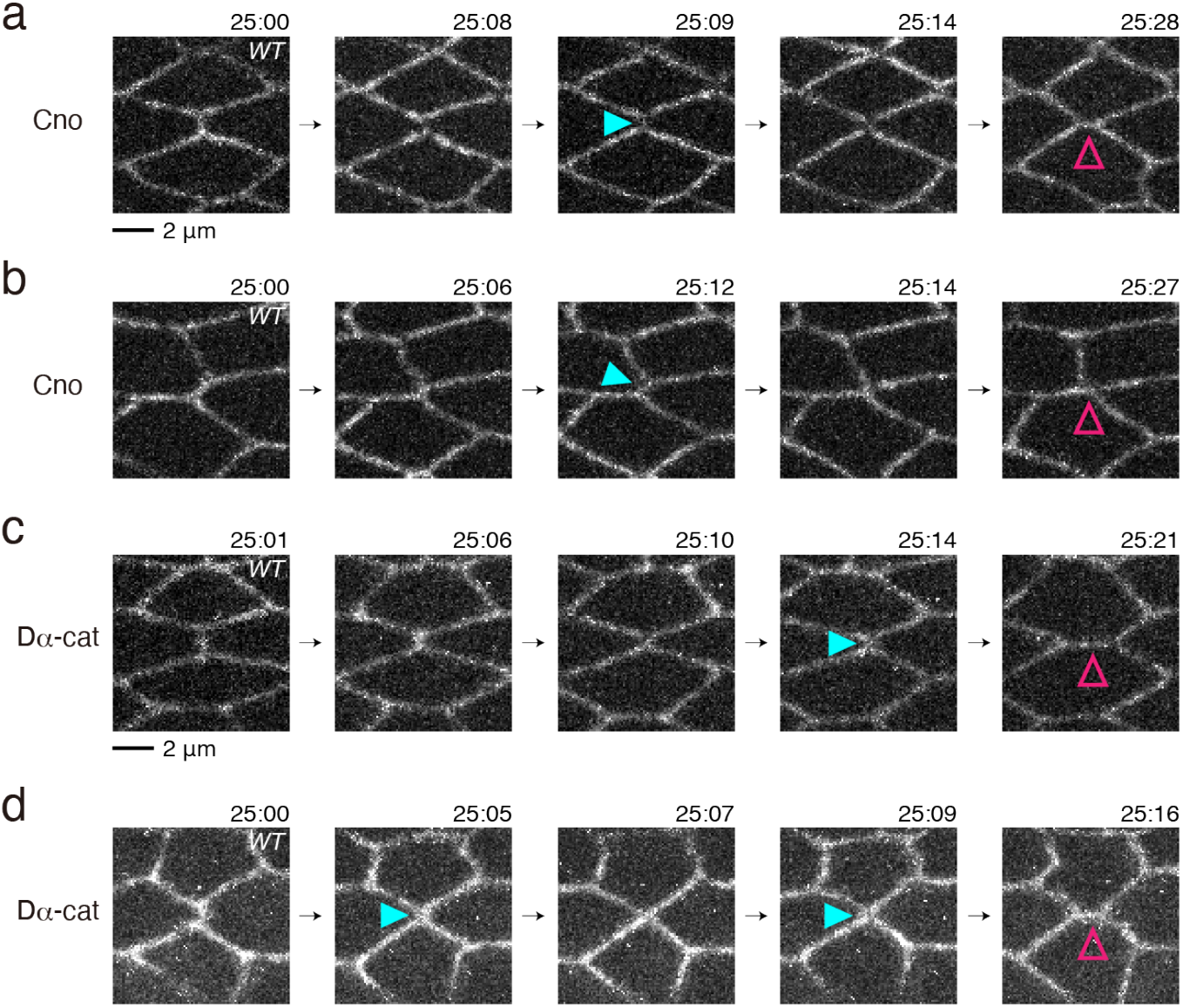
Time-lapse images of Cno and Dα-cat during the junction exchange. We analyzed the distributions of Cno and Dα-cat during the junction exchange in *WT* cells. The Cno and Dα-cat signals were moderately weakened or nearly unchanged during the junction shrinkage and exchange. (a, b) Time-lapse images of Cno-YFP. (c, d) Time-lapse images of Dα-cat-GFP. Scale bars: 2 μm (a, c).

**Supplementary Fig. 6.**
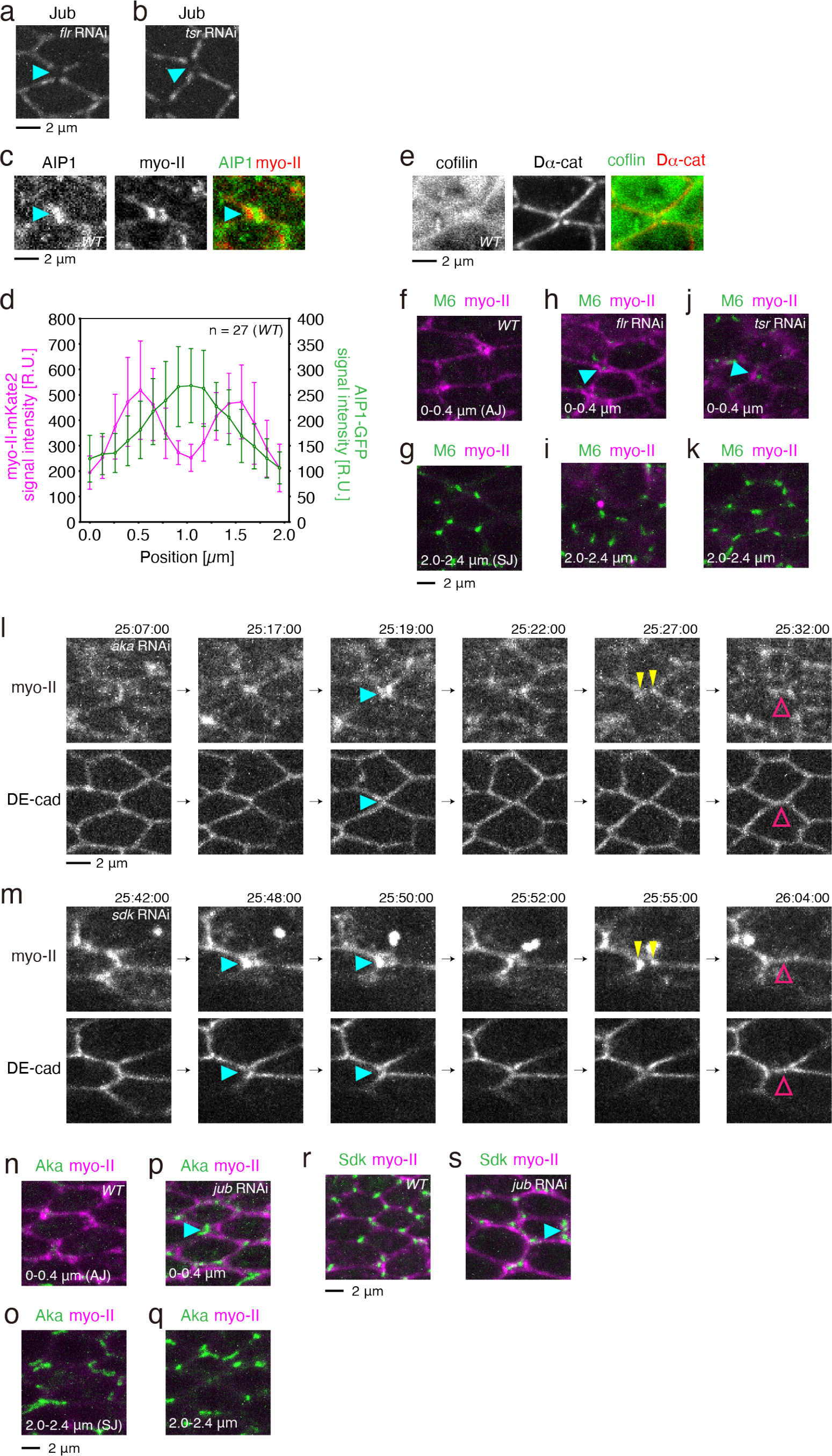
AIP1 and cofilin are required for the correct localization of Jub and M6 during cell rearrangement. We previously showed that actin interacting protein 1 (AIP1) and cofilin retained Cno and Da-cat at the shrinking AP junctions to prevent the precocious formation of the rsMC^16^. Here, by using RNAi against *flare (flr; Drosophila aip1* gene) and *twinstar (tsr; Drosophila cofilin* gene), we confirmed that AIP1 and cofilin are required for the localization of Jub at short junctions. Because *jub* RNAi resulted in a shift in M6 to the AJ plane (Fig. 4a–f), we examined whether Jub is required for localization of other TCJ proteins. Aka shifted to the AJ plane in *jub* RNAi cells as did M6. Sdk localization was largely normal upon *jub* RNAi expression, except that some cells showed ectopic Sdk accumulation along the junction. (a, b) Images of Jub-GFP in wings of the indicated genotypes (a, *flr* RNAi and b, *tsr* RNAi). Images were acquired at 58 h APF at 17°C (a) and 21 h APF at 29°C (b). Blue arrowheads indicate the loss of Jub from AP junctions. (c) Images of AIP1-GFP (gray in left panels, green in right panels) and myo-II-mKate2 (gray in middle panels, red in right panels) at 24.1 h APF. The signal intensity of fluorescent markers along the dashed line is plotted in (d). Arrowheads point to the accumulation of AIP1. In (c) and (e), protein trap lines, in which GFP exons are inserted into the endogenous locus of *flr* and *tsr*, were used, respectively^43,44^. (d) Quantifications of myo-II-mKate2 signal intensity (magenta, left axis) and AIP1-GFP signal intensity (green, right axis) at the rsMC based on time-lapse data captured at 24–26 h APF at 25 °C. The signal intensity of fluorescent markers along the line at each position along the PD axis is plotted. (e) Images of cofilin-GFP (gray in left panel, green in right panel) and Dα-cat-TagRFP (gray in middle panel, red in right panel) at 24 h APF. (f–k) Images of M6-GFP (green) and myo-II-mKate2 (magenta) in *WT* (f, g), *flr* RNAi (h, i), and *tsr* RNAi (j, k) wings. Images along the AJ planes (f, h, j) and more basal planes (g, i, k) are shown as indicated at the bottom. Images were taken at 24 h APF at 25 °C (*WT*), at 58 h APF at 17 °C (*flr* RNAi), and 21 h APF at 29°C (*tsr* RNAi). (l) Time-lapse images of myo-II-mKate2 (upper) and DE-cad-GFP (lower) in the *aka* RNAi wing. Blue, yellow, and magenta arrowheads indicate the detachment of myo-II from the AJ, the separation of the myo-II signals, and the formation of the PD junction, respectively. See Fig. 3c–e for results of the quantification. (m) Time-lapse images of myo-II-mKate2 in the *sdk* RNAi wing. Blue, yellow, and magenta arrowheads indicate the detachment of myo-II from the AJ, the separation of the myo-II signals, and the formation of the PD junction, respectively. (n–q) Images of Aka-GFP (green) and myo-II-mKate2 (magenta) in *WT* (n, o) and *jub* RNAi (p, q) wings at 24 h APF. Images along the AJ planes (n, p) and more basal planes (o, q) are shown as indicated at the bottom. (r, s) Images of Sdk-GFP (green) and myo-II-mKate2 (magenta) in *WT* (r) and *jub* RNAi (s) wings at 24 h APF. Blue arrowhead indicates the ectopic accumulation of Sdk along the junction. Data are presented as the mean ± s.d. (d). Scale bars: 2 μm (a, c, e, g, l, o, r).

## Supplementary Videos

**Supplementary Video 1 myo-II-mKate2 and DE-cad-GFP in *WT* cells.**

Time-lapse images of myo-II-mKate2 (gray in left panels, red in right panels) and DE-cad-GFP (gray in middle panels, green in right panels) in *WT* wing.

Scale bar: 2 μm.

**Supplementary Video 2 Life-Act-GFP and myo-II-mKate2 in *WT* cells.**

Time-lapse images of Life-Act-GFP (gray in left panels, green in right panels) and myo-II-mKate2 (gray in middle panels, red in right panels) in *WT* wing. Lifeact-GFP was clonally expressed.

Scale bar: 2 μm.

**Supplementary Video 3 myo-II-mKate2 and DE-cad-GFP in cells expressing dsRNA against actin-AJ linkers.**

Time-lapse images of myo-II-mKate2 (gray in left panels, red in right panels) and DE-cad-GFP (gray in middle panels, green in right panels) in *jub* RNAi, *eno* RNAi, *a-cat* RNAi, and *vinc* RNAi wings.

Scale bar: 2 μm.

**Supplementary Video 4 Jub-GFP and myo-II-mKate2 in *WT* cells.**

Time-lapse images of Jub-GFP (gray in left panels, green in right panels) and myo-II-mKate2 (gray in middle panels, red in right panels) in *WT* wing.

Scale bar: 2 μm.

**Supplementary Video 5 myo-II-mKate2 and DE-cad-GFP in *m6* RNAi cells.**

Time-lapse images of myo-II-mKate2 (gray in left panels, red in right panels) and DE-cad-GFP (gray in middle panels, green in right panels) in *m6* RNAi wing.

Scale bar: 2 μm.

**Supplementary Video 6 Jub-GFP and myo-II-mKate2 in *m6* RNAi cells.**

Time-lapse images of Jub-GFP (gray in left panels, green in right panels) and myo-II-mKate2 (gray in middle panels, red in right panels) in *m6* RNAi wing.

Scale bar: 2 μm.

**Supplementary Video 7 myo-II-mKate2 and DE-cad-GFP in *pten* RNAi cells.**

Time-lapse images of myo-II-mKate2 (gray in left panels, red in right panels) and DE-cad-GFP (gray in middle panels, green in right panels) in *pten* RNAi wing.

Scale bar: 2 μm.

